# Embryonic hyperglycemia perturbs the development of specific retinal cell types, including photoreceptors

**DOI:** 10.1101/2021.05.02.442186

**Authors:** Kayla F. Titialii-Torres, Ann C. Morris

## Abstract

Chronic hyperglycemia has been linked to various long-term metabolic disruptions in adults, such as neuropathy, nephropathy, and diabetic retinopathy. According to the 2020 National Diabetes Statistics Report, 10.5% of the US population has diabetes and may be susceptible to long-term complications if blood glucose is not tightly regulated. Further, in 2018, 7.6% of US pregnancies were affected by gestational diabetes, with an average of 1-14% annually [1]. During pregnancy, glucose can pass through the placental barrier, and plays an important role in fetal development and survival. However, excess maternal glucose can also result in diabetic embryopathy. While many studies have examined the teratogenic effects of maternal diabetes on fetal heart development, little is known about the consequences of maternal hyperglycemia on the development of the embryonic retina. To address this question, we investigated retinal cell type differentiation and survival in both a genetic and nutritional model of embryonic hyperglycemia in zebrafish. Strikingly, we found that hyperglycemic larvae displayed a significant reduction in rod and cone photoreceptors and horizontal cells, whereas other retinal neurons were not affected. We also observed signs of reactive gliosis in the retinal Müller cells, as well as increased reactive oxygen species (ROS) production in hyperglycemic retinas. Hyperglycemic larvae displayed altered expression of metabolism related genes and had a slower optokinetic response than normoglycemic larvae, indicating altered visual function. Further analysis of early events in retinogenesis revealed a delay in retinal cell differentiation at 48 hpf in hyperglycemic embryos, that coincided with an increase in reactive oxygen species. Taken together, our results suggest that embryonic hyperglycemia results in abnormal retinal cell development via altered timing of retinal cell differentiation and ROS production, which is accompanied by visual defects. As the population with diabetes continues to grow, it is imperative to pinpoint the effects of embryonic hyperglycemia on retinal development. Further studies using zebrafish models of hyperglycemia will allow us to understand the molecular mechanisms underlying these effects, which could aid in the development of therapeutic strategies.

## Introduction

Diabetes is a growing epidemic, affecting 34.2 million people in the US in 2018 [2]. High blood sugar (hyperglycemia) is the primary symptom of diabetes and when it becomes a recurring state, various complications are likely to arise which affect tissues all over the body [3]. Complications include neuropathies (nerve damage), nephropathy (kidney disease), stroke, and retinopathy which can cause progressive blindness [4]. While these complications are primarily documented in adults who have experienced recurring hyperglycemia over many years, hyperglycemia during pregnancy carries its own set of complications which can have long lasting effects on the offspring [5]–[7].

During pregnancy, glucose passes through the placental barrier from the mother to fetus; this maternally supplied glucose is necessary for fetal development but in excess it can also result in embryonic hyperglycemia. Maternal hyperglycemia may come from existing diabetes prior to pregnancy and/or increased insulin resistance developed during pregnancy to allow for increased glucose to pass through the placenta, which is essential for stimulating fetal insulin production to aid in growth [8]. Excess insulin resistance can lead to gestational diabetes which is not diagnosed until 24 weeks into pregnancy (and occurs in nearly 10% of US pregnancies). The type and severity of hyperglycemia related complication varies greatly, depending on when embryonic hyperglycemia occurs. The most prominently studied complication of embryonic hyperglycemia is heart malformation or congenital heart defect (CHD). A wide array of phenotypes are associated with CHD, and it is considered the most common birth defect associated with diabetic embryopathy [9]. However, other developing tissues and organs are also vulnerable to the effects of embryonic hyperglycemia, and these have been less well studied. Given the strong connection between diabetes and retinal degeneration leading to progressive blindness in adults, there is a critical need to study the effects of hyperglycemia on the developing retina during embryogenesis, using an animal model where the developing eye is easily accessible.

Zebrafish have recently become a favorable model for studying hyperglycemia and diabetes due to their relatively easy maintenance, high fecundity, and manipulatable environment.

Diabetes and hyperglycemia can be induced via ablation of pancreatic beta cells through streptozotocin injection [10] or whole-body immersion in glucose dense fish water [11]. Using the immersion technique with adult zebrafish, it was demonstrated that recurring hyperglycemia resulted in a reduction in the number of cone photoreceptors in the retina, with remaining photoreceptors displaying an abnormal morphology, including shortened outer segments [12] as well as abnormal electroretinogram responses [13]. Recent studies utilizing a genetic mutant to induce hyperglycemia (*pdx1*^*-/-*^) showed that recurring hyperglycemia in adulthood resulted in photoreceptor degeneration, defective visual responses [14], and increased retinal angiogenesis, a hallmark of diabetic retinopathy [15]. While these studies show the utility of zebrafish to study the ocular complications of hyperglycemia in adults, there has been less work on the effects of hyperglycemia during embryonic and larval retinal development. One recent study suggested that exposure to very high levels of exogenous glucose causes a decrease in retinal ganglion cells and Müller glia as well as an increase in vasculature leakage in zebrafish larvae [16], providing evidence that embryonic hyperglycemia has deleterious effects on retinal development. However, the consequences of embryonic hyperglycemia specifically on photoreceptor development have not been closely examined. Given that there is mounting evidence that photoreceptors, which are highly metabolically demanding cells, are major contributors to the progression of diabetic retinopathy, there is a pressing need to study how hyperglycemia may impact photoreceptors during embryonic development.

In this study, we specifically explored the consequences of hyperglycemia on cell type differentiation in the developing zebrafish retina, using two complementary approaches. To model chronic hyperglycemia via lack of insulin production, similar to what is observed in type I diabetes, we utilized *pdx1*^*-/-*^ zebrafish, which possess a null mutation in a gene necessary for beta cell development, and therefore cannot produce insulin [17]. A recent study showed *pdx1*^-/-^ zebrafish larvae have microvascular changes in the ocular hyaloid vasculature at 6 days post fertilization (dpf) [15], however an examination of photoreceptor and retinal development has not yet been reported for this mutant. To model hyperglycemia that is representative of type II diabetes, we developed a nutritional model, in which zebrafish embryos are exposed to exogenous glucose and dexamethasone from 10 hours post fertilization (hpf), just prior to optic vesicle evagination, until 5 dpf, when retinal development is largely complete. Dexamethasone is a synthetic glucocorticoid that is used in combination with glucose to elevate whole body glucose due to its ability to stimulate gluconeogenesis and disrupt glucose transport [18]. In the context of embryonic development, dexamethasone is often given to pregnant women who are at risk of preterm birth to aid in fetal lung development, and it is known that low birth weight and preterm babies are highly susceptible to hyperglycemia [19].

Here we report that, in both genetic and nutritional models of embryonic hyperglycemia, rod and cone photoreceptor cells are significantly decreased in number, and retinal oxidative stress is increased. Notably, embryonic hyperglycemia was associated with abnormal visual behavior at 5 dpf. Additionally, the timing of retinal progenitor differentiation was altered in hyperglycemic larvae, and cone photoreceptor number remained lower the controls even after a return to normoglycemic conditions. These findings provide evidence that embryonic hyperglycemia impedes proper retinal development, leading to short-term, and potentially long-term, visual defects.

## Results

### Hyperglycemia is detectable in pdx1 mutant larvae at 5 dpf

Prior studies of two different *pdx1* mutant zebrafish lines have demonstrated that at 5 dpf, mutant larvae have elevated whole body glucose, aberrant hyaloid vasculature, and later grow to be significantly smaller compared to their wildtype and heterozygous siblings [15], [17]. Using the *pdx1*^sa280^ mutant line described by Kimmel et al, we found that *pdx1* mutants displayed elevated whole-body glucose levels as early as 4 dpf (data not shown) and confirmed that they have significantly elevated whole-body glucose at 5 dpf (Fig. S1A). We note that whole-body glucose was measured rather than blood glucose because the total blood volume of a 5 dpf zebrafish larvae is less than a microliter [20]. We measured *pdx1* mutant eye size and found that it was proportional to their body size when compared with wild type larvae, indicating that the mutation and accompanying hyperglycemia does not cause microphthalmia (Fig. S1B-D). However, further analysis at the cellular level revealed interesting abnormalities in the developing retina.

### Pdx1 mutant larvae have reduced numbers of photoreceptors

To determine whether hyperglycemia impacts photoreceptor development, we crossed heterozygous *pdx1*^*sa280*^ adults onto the XOPS:GFP transgenic background, in which rod photoreceptors are fluorescently labeled [21]. *Pdx1*^*+/-*^ ;XOPS:GFP^+/-^ and *pdx1*^*+/-*^ adults were in-crossed to assess rod photoreceptor number and morphology. We observed a significant decrease in rod photoreceptors of *pdx1*^*-/-*^ larvae at 5 dpf compared to wildtype and heterozygous siblings (Fig. 1F). Looking closely at the rod photoreceptors in the ventral retina, the outer segments appeared to be much shorter and thinner (Fig. 1E’ arrowhead) when compared to wildtype or heterozygous retinas (Fig. 1D’). The dorsal retina also contained decreased numbers of rod photoreceptors in mutants (Fig. 1E) compared to wildtype (Fig. 1D). Similarly, red/green cone photoreceptors in the *pdx1*^*-/-*^ retina, labeled by the Zpr1 antibody, displayed a decrease in number (Fig. 1B) compared to wildtype retinas (Fig. 1A). In the ventral region cones possessed stunted outer segments and appeared to have a thinner cell body than those in wildtype larvae (Fig. 1B’).

**Fig. 1.**
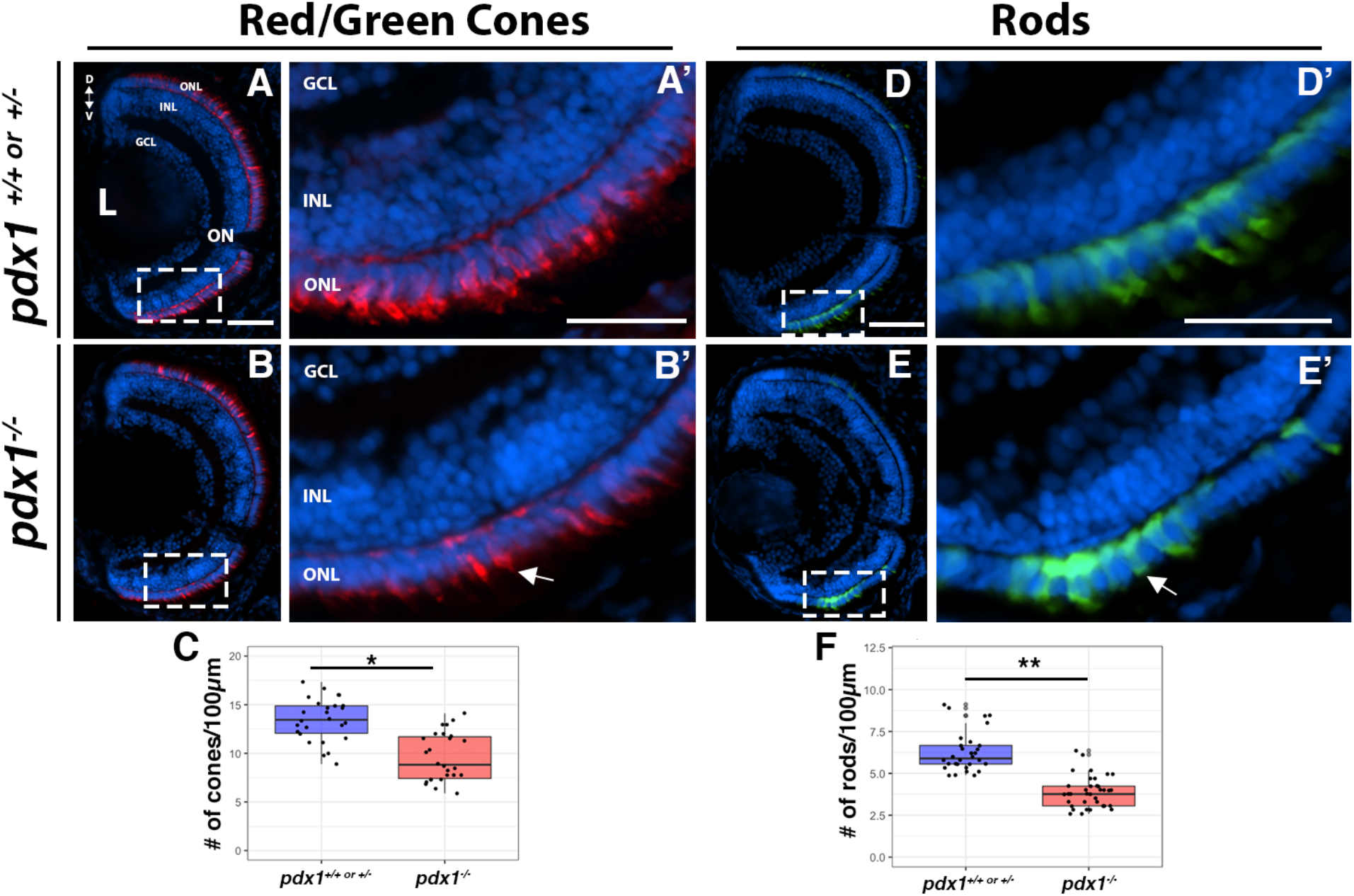
Photoreceptors display reduced numbers and abnormal morphology in *pdx1*^-/-^ larvae at 5 dpf. Larvae from *pdx1*^*+/-*^ incrosses were genotyped and sectioned at 5 dpf, then immunostained and imaged to visualize photoreceptors. Red/green double cones were visualized with the Zpr1 antibody (A-B) and showed a significant reduction in number in *pdx1*^-/-^ larvae (C). The ventral portion of the retina shows a stark difference in morphology and spacing of red/green double cones in *pdx1*^-/-^ larvae (B’). Rod photoreceptors were visualized using the XOPS:GFP transgene (D-E) and showed a significant reduction in number in *pdx1*^-/-^ larvae (F). The ventral portion of the retina is emphasized in D’ and E’, showing the difference in cell morphology. D, Dorsal; V, Ventral; ONL, Outer Nuclear Layer; INL, Inner Nuclear Layer; GCL, Ganglion Cell Layer; L, Lens. Scale bars: 50µM (A, B) and 20µM (A’, B’). * indicates p<0.05; ** indicates p<0.01

Quantification revealed a significant decrease in both rod and cone photoreceptors in *pdx1* mutants at 5 dpf (Fig. 1C). The average number of red/green double cone photoreceptors was reduced by over 20% which was particularly striking in the ventral portion of the retina (average of 13/100µm in WT or het vs. 9 in *pdx1*^*-/-*^ mutants), and the average number of rods was decreased by 45% (average of 5.5/100µm in wildtype or het vs. 3 in *pdx1*^*-/-*^ mutants). Taken together, we conclude that hyperglycemic *pdx1* mutants have significantly less photoreceptors than normoglycemic larvae of the same age, and the photoreceptors that are present in *pdx1* mutant retinas display an abnormal morphology.

### Induction of hyperglycemia in developing zebrafish via glucose and dexamethasone exposure

While the *pdx1* mutant provides insight into hyperglycemic phenotypes resulting from a genetic defect in insulin producing cells, we also wanted to determine whether embryonic and larval hyperglycemia induced by altered nutrient uptake had similar effects on retinal development. With this approach, we can model the effects of a direct, in utero, exposure to elevated glucose that may occur in cases of maternal diabetes, and which involves excess glucose flow across the placenta [22].To that end, we submerged zebrafish embryos in fish water containing glucose +/-dexamethasone from 10 hpf (just prior to optic vesicle evagination from the forebrain) until 5 dpf (when retinal development is largely complete). To avoid nonspecific effects of high glucose on embryo development, we conducted a series of dose responses for glucose concentrations and selected 50mM as it was the lowest concentration which resulted in whole body glucose elevation. This is consistent with a previous study, which used 55mM glucose treatments to induce hyperglycemia and which resulted in abnormal vasculature development [23]. Other studies involving a glucose submersion technique with zebrafish to induce hyperglycemia have used higher glucose concentrations that ranged from 110mM to 277mM [13], [16].

Treatment of zebrafish embryos with glucose alone produced a significant increase in whole body glucose in comparison to untreated embryos and to embryos exposed to mannitol (osmolarity control; Fig. 2B). While whole body glucose was significantly elevated in response to glucose treatment, we found that exposure to glucose alone resulted in highly variable levels of hyperglycemia. To combat this, we added dexamethasone to the glucose treatment. Dexamethasone is a synthetic glucocorticoid that has been shown to disrupt glucose transport into cells, preventing proper breakdown of glucose and increased free glucose in the blood [24]. Dexamethasone is also frequently given antenatally to facilitate fetal lung maturation [25]. Our results show the combination of glucose and dexamethasone treatment provides a much tighter range of significantly elevated whole-body glucose values (Fig. 2B). Additionally, dexamethasone treatment alone did not significantly affect whole-body glucose in comparison to untreated and mannitol controls (Fig. 2B).

**Fig. 2.**
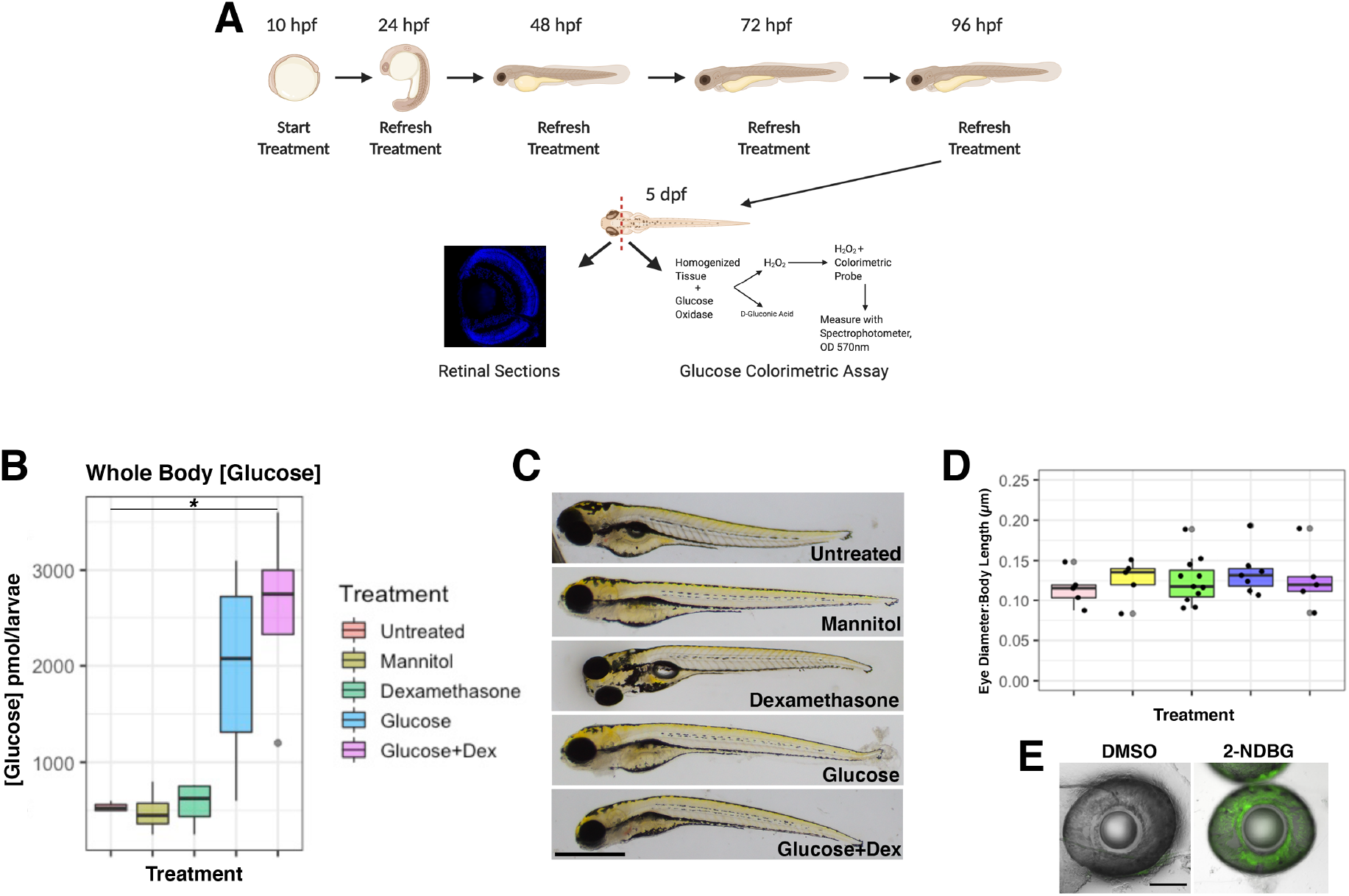
Diagram of methodology to induce hyperglycemia in embryos and larvae (A). Embryos were dechorionated and randomly assorted into groups of 25, then submerged in one of the following treatments: untreated, 50mM mannitol, 10µM dexamethasone, 50mM glucose, 50mM glucose + 10µM dexamethasone (glucose+dex). Beginning at 24 hpf, each treatment was refreshed by replacing with freshly made solutions every 24 hours until 5 dpf. At 5dpf, larval heads were used for cryosectioning the retina, while the rest of the body was used for a glucose colorimetric assay (A). Whole body glucose concentration was elevated in glucose and glucose+dex treated larvae compared to untreated, mannitol, and dexamethasone controls (B). Whole bodies of 5 dpf treated larvae showed no significant differences in eye morphology (C) in relation to body size across treatments (D). Green fluorescence was observed throughout the eye in 2-NDBG treated larvae while none was observed in the eyes of the DMSO control (E). Scale bar: 500µM. * indicates p<0.05; ** indicates p<0.01*

When comparing the gross morphology of larvae from the different treatment groups, we observed some variation in body length as well as yolk and eye size. However, when eye size was normalized to body size, there was no significant difference in ocular proportions across groups, indicating that our experimental conditions do not cause microphthalmia (Fig. 2C-D). Moreover, using a fluorescent glucose analog (2-NDBG) we confirmed that exposure to exogenous glucose leads to glucose uptake in the eye (Fig. 2E). Taken together, our results show that a combination of glucose and dexamethasone exposure reliably produces hyperglycemia in zebrafish larvae. From this point forward we will mostly present data from the mannitol and glucose+dex treatment groups.

### Decreased retinal photoreceptors in a nutritional model of embryonic hyperglycemia

With the establishment of a nutritional model to compare to the *pdx1* mutant, we conducted similar analyses on photoreceptor development. Progeny of XOPS:GFP and TαC:eGFP (a transgenic line that fluorescently labels all cone photoreceptors;[26]) adult in-crosses were used in treatments to assess photoreceptor number and morphology. Similar to the *pdx1* mutant, we observed a significant decrease in both rod and cone photoreceptors of hyperglycemic larvae at 5 dpf (Fig. 3C, F). Cone photoreceptors were decreased by 23% in glucose treated and 28% in glucose+dex treated retinas compared to those treated with the mannitol control (Fig. 3C), whereas rods were decreased by 34% in glucose treated and 45% in glucose+dex treated retinas (Fig. 3F). Compared to *pdx1* mutants, the decrease in cones was larger in the nutritional model, which may be because the TαC:eGFP transgene used for these experiments labels all cone photoreceptor subtypes (red, green, blue, and UV), whereas only red/green cones were detected by the Zpr1 antibody used with the *pdx1* larvae. In contrast, the reduction in rods was comparable in the glucose+dex treated larvae and *pdx1* mutant larvae. Confocal microscopy of whole eyes from control and hyperglycemic larvae revealed a large decrease in the number of rod photoreceptors in hyperglycemic compared to mannitol treated larvae (Fig. 3D, E). In addition, the confocal images showed that, similar to the *pdx1* mutant retinas, the outer segments of both rod and cone photoreceptors appeared stunted compared to controls, with wider inner segments (Fig. 3 B’, E’). Taken together, our results show that embryonic hyperglycemia induced by exogenous glucose results in a reduction of photoreceptors, similar to what is noted in *pdx1* mutants, further supporting a significant impact of embryonic hyperglycemia on photoreceptor development.

**Fig. 3.**
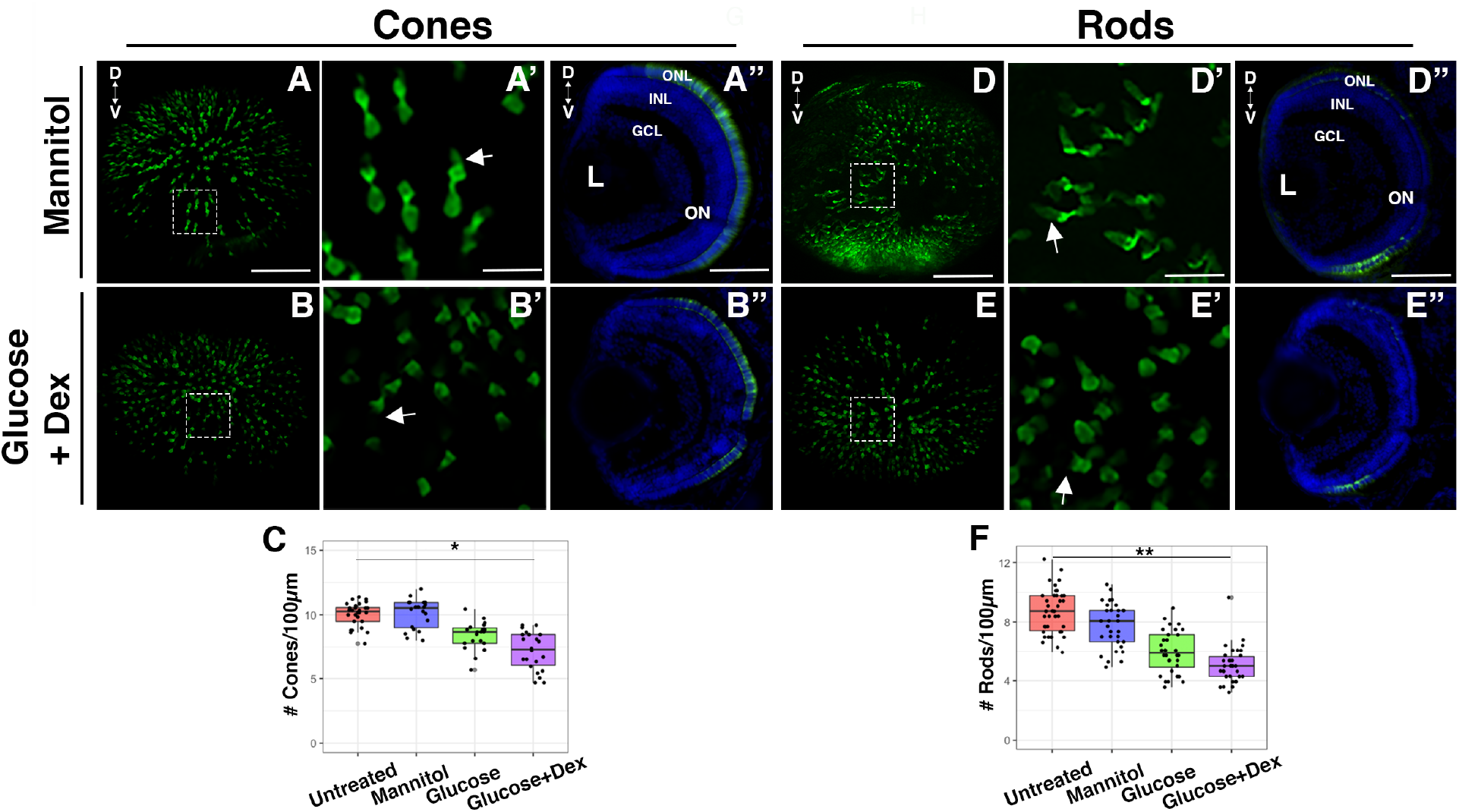
Photoreceptors display reduced number and abnormal morphology in glucose + dex treated larvae at 5 dpf. Treated larvae were collected at 5 dpf, then whole eyes were dissected and imaged by confocal microscopy, or larvae were cryosectioned to visualize retinal photoreceptors. Cone photoreceptors were visualized using the TαC:eGFP trangenic line (A-B”) and rod photoreceptors were visualized using the XOPS:GFP transgenic line (D-E”). Confocal z-stacks of whole eyes (A, B) and selected regions of those stacks reveal long outer segments in mannitol treated embryos (A’, arrow) and truncated outer segments glucose+dex treated larvae (B’, arrow). Cone photoreceptor quantification from retinal sections showed a significant reduction in cone photoreceptors in glucose+dex treated larvae compared to controls (C). Confocal stacks of the XOPS:GFP treated larvae (D, E) revealed large patches of reduced or missing rod photoreceptors in addition to truncated outer segments in glucose + dex treated (E’, arrow) compared to mannitol treated larvae (D’, arrow). Retinal sections confirmed the significant reduction in number of rod photoreceptors (F) in glucose+dex treated larvae (E”) compared to controls (D”). D, Dorsal; V, Ventral; ONL, Outer Nuclear Layer; INL, Inner Nuclear Layer; GCL, Ganglion Cell Layer; L, Lens. Scale bars: 50µM (A, A”, D, D”) and 10µM (A’, D’). * indicates p<0.05; ** indicates p<0.01*

### Hyperglycemic larvae exhibit visual defects

Our results from both models indicate that embryonic hyperglycemia causes photoreceptor defects in the larval retina, specifically in overall number and in outer segment morphology. To further assess outer segments, we used the Zpr3 antibody, which labels the outer segments of rods and red/green double cones [27]. The results confirmed that rods and cones from hyperglycemic larvae have shorter outer segments compared to wild type larvae at 5 dpf (Fig. 4A-C). The purpose of outer segments is to capture light and, via the phototransduction cascade, convert it to an electrical signal to be sent through the retina to the brain [28]. Both *pdx1* mutant and glucose+dex treated larvae displayed abnormally short photoreceptor outer segments. Without full, elongated outer segments, we hypothesized that hyperglycemic larvae may experience subtle visual defects [29]. Therefore, we performed an optokinetic response (OKR) assay to measure larval visual response. Comparison of the number of ocular saccades per minute across *pdx1* genotypes revealed a significant reduction of eye movements in the *pdx1* mutants compared to their wildtype and heterozygote siblings (Fig. 4C). Similarly, glucose+dex treated larvae also showed a significant decrease in OKR performance at 5 dpf compared to untreated, mannitol treated, and dexamethasone treated larvae (Fig. 4F). Glucose treated larvae showed a wide range in response, reflective of the variability in their photoreceptor number and morphology. Together, these results show that the photoreceptor defects observed in hyperglycemic larvae are associated with reduced visual responses. The similarity in phenotype between *pdx1* mutants and glucose+dex treated larvae, with respect to photoreceptor number, photoreceptor morphology, and visual acuity, indicates that these phenotypes are due to hyperglycemia. Therefore, we next wanted to delve into the mechanisms by which these phenotypes arise.

**Fig. 4:**
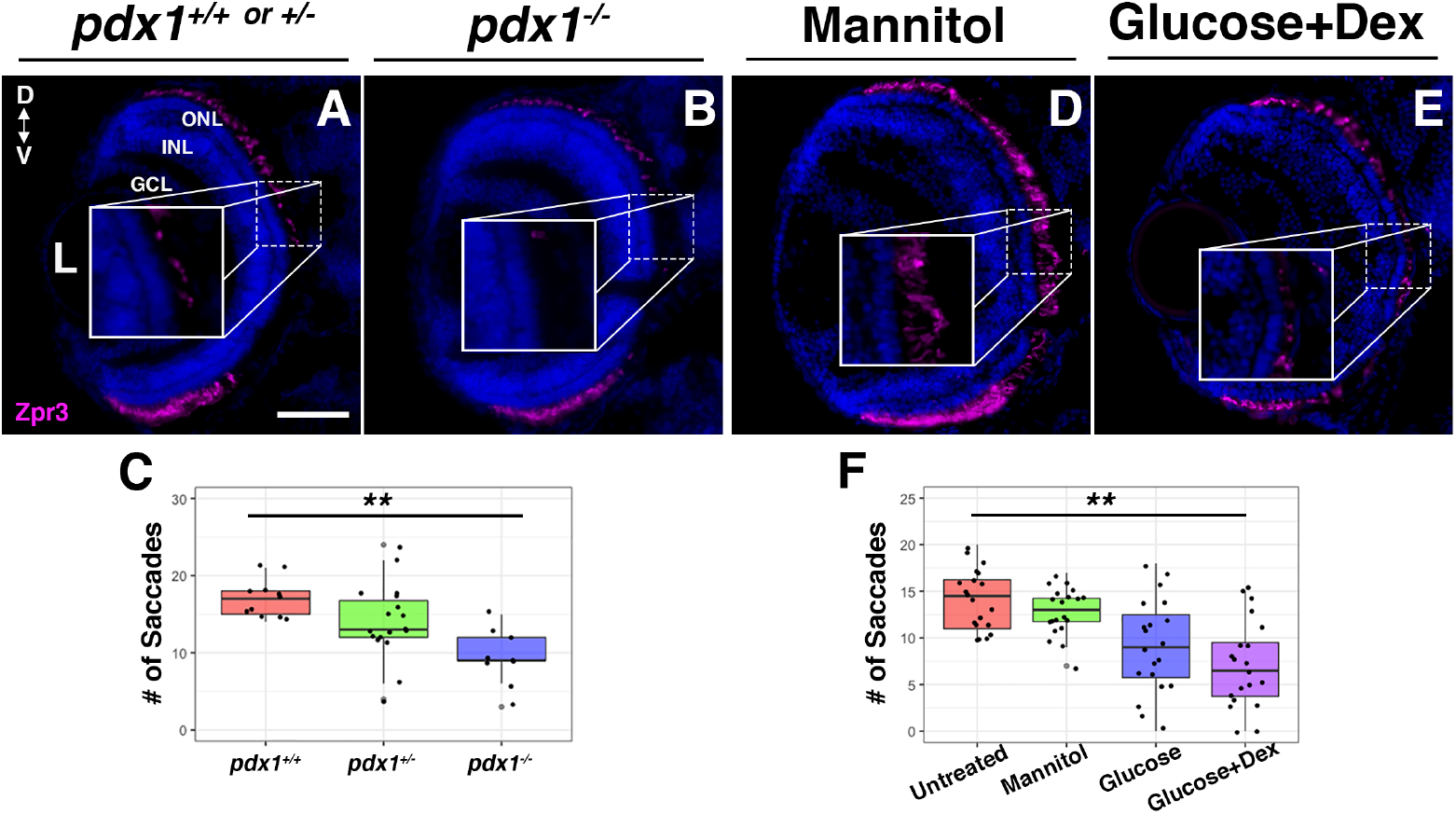
Hyperglycemic larvae have truncated outer segments and subtle visual defects. Pdx1 larvae were genotyped, then both Pdx1 and treated larvae were sectioned at 5 dpf, immunostained and imaged to visualize outer segments. Immunostaining for outer segments with Zpr3 showed a decrease in density of outer segments in both hyperglycemic models (B, E), with a particularly striking difference in the ventral portion of the retina compared to controls (A, D). Optokinetic Response Test was also conducted on larvae at 5 dpf by placing the larvae inside a rotating drum with alternating black and white stripes, then quantifying the number of saccadic eye movements (saccades). Hyperglycemic larvae showed a decrease in saccades compared to controls (C, F). D, Dorsal; V, Ventral; ONL, Outer Nuclear Layer; INL, Inner Nuclear Layer; GCL, Ganglion Cell Layer; L, Lens. Scale bars: 50µm (A). ** indicates p<0.01

### Apoptosis is modestly elevated in hyperglycemic larval retinas

Hyperglycemia has been shown to induce cell death via apoptosis and autophagy as well as to increase susceptibility to necrosis [30]–[32]. Therefore, we wanted to quantify and compare programmed cell death in control and hyperglycemic larvae. Using the TUNEL assay we detected apoptotic cells in retinal sections at 5 dpf. We found a significant increase in apoptotic cells with both the genetic and nutritional models of hyperglycemia (Fig. S2C, F), primarily in the INL and to a lesser extent in the ONL (Fig. S2B, E). Looking at additional timepoints leading up to 5 dpf, we found cell death began to significantly increase at 4 dpf (Fig. S3Y), which aligns with the first time point at which we see elevation in whole body glucose in *pdx1* mutants and glucose+dex treated larvae. Although there was a significant increase in apoptosis in hyperglycemic retinas, the total number of TUNEL+ cells per retinal section was low across all treatments, genotypes, and timepoints. Interestingly, most of the TUNEL+ cells were located in the central portion of the inner nuclear layer, coinciding with the location of Müller glia cell bodies. To follow up on this, we assessed Müller glia morphology and number in hyperglycemic retinas.

### Hyperglycemic larval retinas exhibit signs of reactive gliosis

Sections of control and hyperglycemic GFAP:GFP (a transgenic line in which Müller glia cells are GFP tagged) larvae revealed an increase in average number of Müller glia in glucose treated larvae (+17%) at 5 dpf and a slight increase in the glucose+dex treated larvae (+11%) compared to untreated and mannitol treated controls (Fig. 5G). Looking at the distribution of Müller glia within the inner nuclear layer, glucose+dex treated larvae displayed an irregular localization of Müller cell bodies (Fig. 5F) compared to the linear pattern of cell bodies in mannitol and glucose treated retinas (Fig. 5B, 5D). Moreover, whereas Müller glial cell bodies possessed the expected compact polygonal shape in mannitol and glucose treated retinal sections, the Müller glial cell bodies of glucose+dex treated retinas appeared swollen, twisted, or heart-shaped, which could represent signs of reactive gliosis (Fig. 5F). For a better understanding of Müller glia morphology, we utilized whole-mount confocal microscopy to visualize Müller glia in 3-D space. We found the increase of Müller glia in glucose treated larvae was evident throughout the eye (Fig. 5C, C’), while the shape and size of Müller glia were comparable to mannitol treated retinas (Fig. 5A, A’). Whole mount imaging of glucose+dex treated larvae revealed a variety of abnormal cell body shapes throughout the eye (Fig. 5E, E’) which were also larger than glucose and mannitol treated. Together, these results indicate a gliotic response in the glucose+dex treated retinas at 5 dpf. We also imaged sections of *pdx1* larvae at 5 dpf that were crossed onto the GFAP:GFP background. Unlike glucose and glucose+dex treated larvae, *pdx1* mutants did not exhibit an increase in the number of Müller glia (Fig. 5K). Interestingly though, we did find that Müller glia cell bodies in *pdx1*^*-/-*^ retinas were significantly larger than their wildtype and heterozygous counterparts (Fig. 5L) and displayed a variety of abnormal shapes similar to glucose+dex treated larvae, with some disorganization in patterning (Fig. 5J). Taken together, our results suggest that embryonic hyperglycemia induces a gliotic response within the larval retina. Furthermore, exogenous exposure to glucose causes a slight increase in Müller glial number that is not observed in *pdx1* mutant retinas, suggesting that this effect may be due to elevated levels of insulin signaling in our nutritional model.

**Fig. 5:**
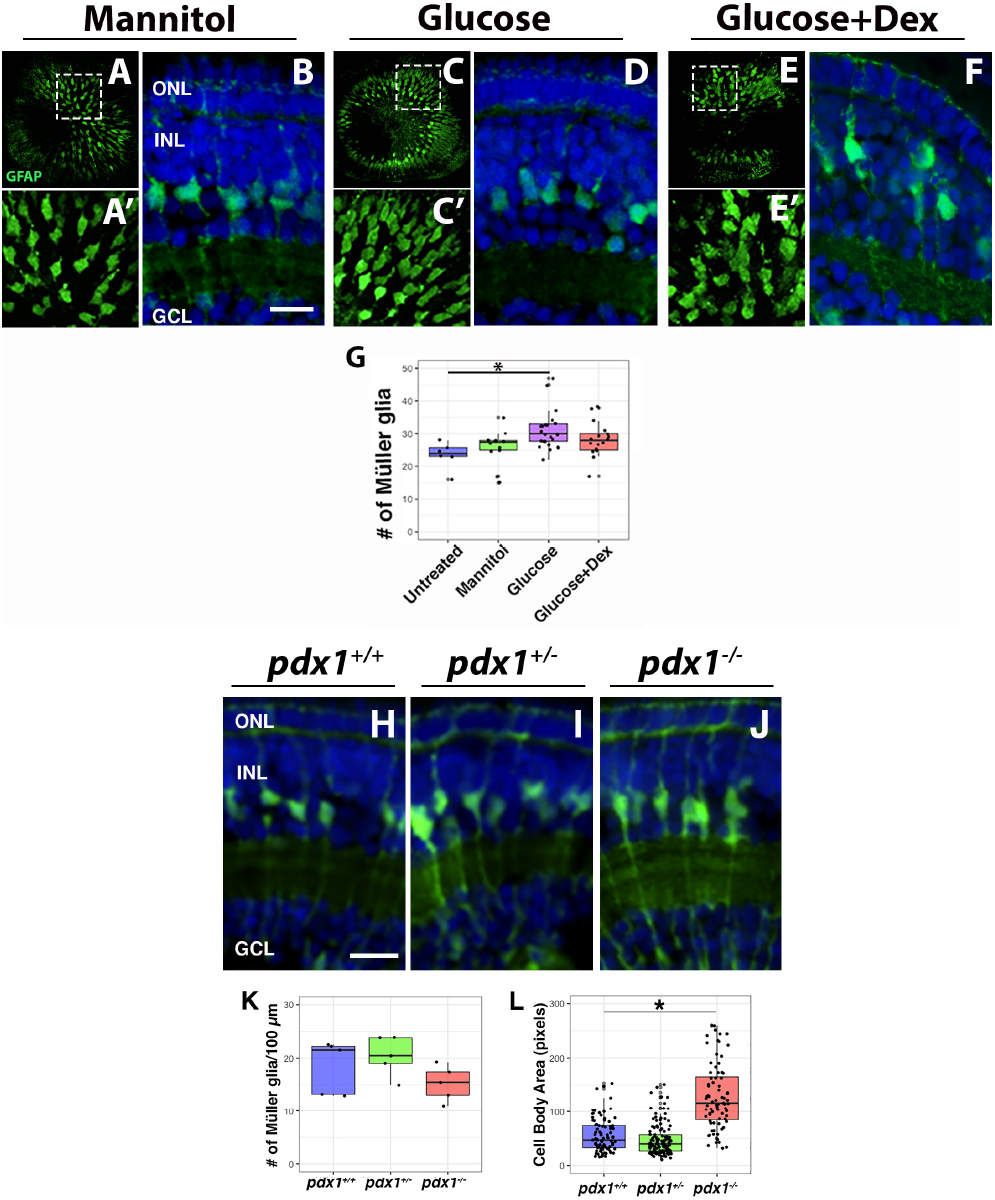
Signs of reactive gliosis in hyperglycemic larvae. Wildtype and *pdx1*^*+/-*^ adults were crossed onto the GFAP:GFP transgenic background to visualize Müller glia. At 5 dpf, whole mount confocal images (A, B, C) of treated wildtype larvae showed a slight increase in the number of Müller glia in glucose treated retinas (C). Glucose+dex treated larvae displayed large, irregular shaped nuclei (E’). Retinal sections confirmed the increased number of Müller glia in glucose treated larvae (D) and the enlarged nuclei in glucose+dex treated retinas (F). Retinal sections of *pdx1* mutant larvae crossed onto the GFAP:GFP transgenic background at 5 dpf (H-J) showed no difference in number of Müller glia (K) but a significant increase in the size of Müller glia cell bodies (L). ONL, Outer Nuclear Layer; INL, Inner Nuclear Layer; GCL, Ganglion Cell Layer. Scale bars: 10µm. * indicates p<0.05

### Horizontal cells are reduced in hyperglycemic larvae while bipolar, amacrine, and ganglion cell numbers are unchanged

To determine whether embryonic hyperglycemia resulted in altered numbers of other retinal neurons, we performed immunohistochemistry with antibodies that label various retinal cell types. Horizontal cells are located in the outermost part of the inner nuclear layer, and synapse directly to photoreceptors. We detected horizontal cells with the Prox1 antibody which also labels progenitor cells in the inner nuclear layer. Horizontal cells were identified by their location and distinct oblong shape. There was a 30% decrease in the number of Prox1+ horizontal cells per 100µm in *pdx1* mutant retinas compared with wildtype (Fig. S4A-C). In the nutritional model, both glucose and glucose+dex treated embryos displayed a 40% and 50% decrease, respectively, in horizontal cell number (Fig S5). Bipolar cells, which also synapse directly with photoreceptors and transfer the visual signal from photoreceptors to ganglion cells ([33]), did not show a difference in average number per 100µm when comparing wildtype to *pdx1* mutants (Fig. S4D-F) or in glucose and glucose+dex treated larvae compared to controls (Fig. S5D-F). Next, we quantified the number of ganglion and amacrine cells, using the HuC/D antibody. There was no significant difference in amacrine cell number across genotypes (Fig. S4G-I) or treatments (Fig. S5G-I). Similarly, we found no significant differences across genotypes or treatments in the number of retinal ganglion cells (Fig. S4 G-I and S5 G-I). These results suggest that abnormal retinal cell type differentiation in hyperglycemic larvae is limited to photoreceptors and horizontal cells, with the remaining inner retinal neurons developing in normal numbers.

### Reactive oxygen species production is increased in hyperglycemic larvae via dysregulation of glucose metabolism-related pathways

Hyperglycemia has been shown to induce increased levels of reactive oxygen species (ROS), unstable oxygens that cause DNA damage and cell death [34]. A recent study showed ROS production is a key component of retinal development through regulation of a switch between cellular proliferation and differentiation [35]. Further, studies in adult mice and retinal explants have shown that recurring hyperglycemia leads to an increase in ROS production, specifically in photoreceptors [36], [37]. To determine whether ROS production was increased in the retinas of hyperglycemic larval zebrafish we used Mitosox, an in-vivo probe for superoxides. We found increased ROS production in both glucose and glucose+dex treated embryos at 48hpf, a key window of photoreceptor differentiation (Fig. 6D, E). At 5 dpf we noted a significant increase in ROS in pdx1 mutants and glucose+dex (Fig. S6B, D). While ROS probe signal was observed at various places throughout the head, it was especially prominent in the eyes of hyperglycemic larvae (Fig. 6D, E and S6D). Retinal sections at 5dpf showed the ROS in the retina to be specifically located among the outer segments of photoreceptors (Fig. S6F, H, arrows). In contrast, wildtype and untreated larvae had long, straight outer segments which did not show ROS probe colocalization (Fig. S6E, G).

**Fig. 6:**
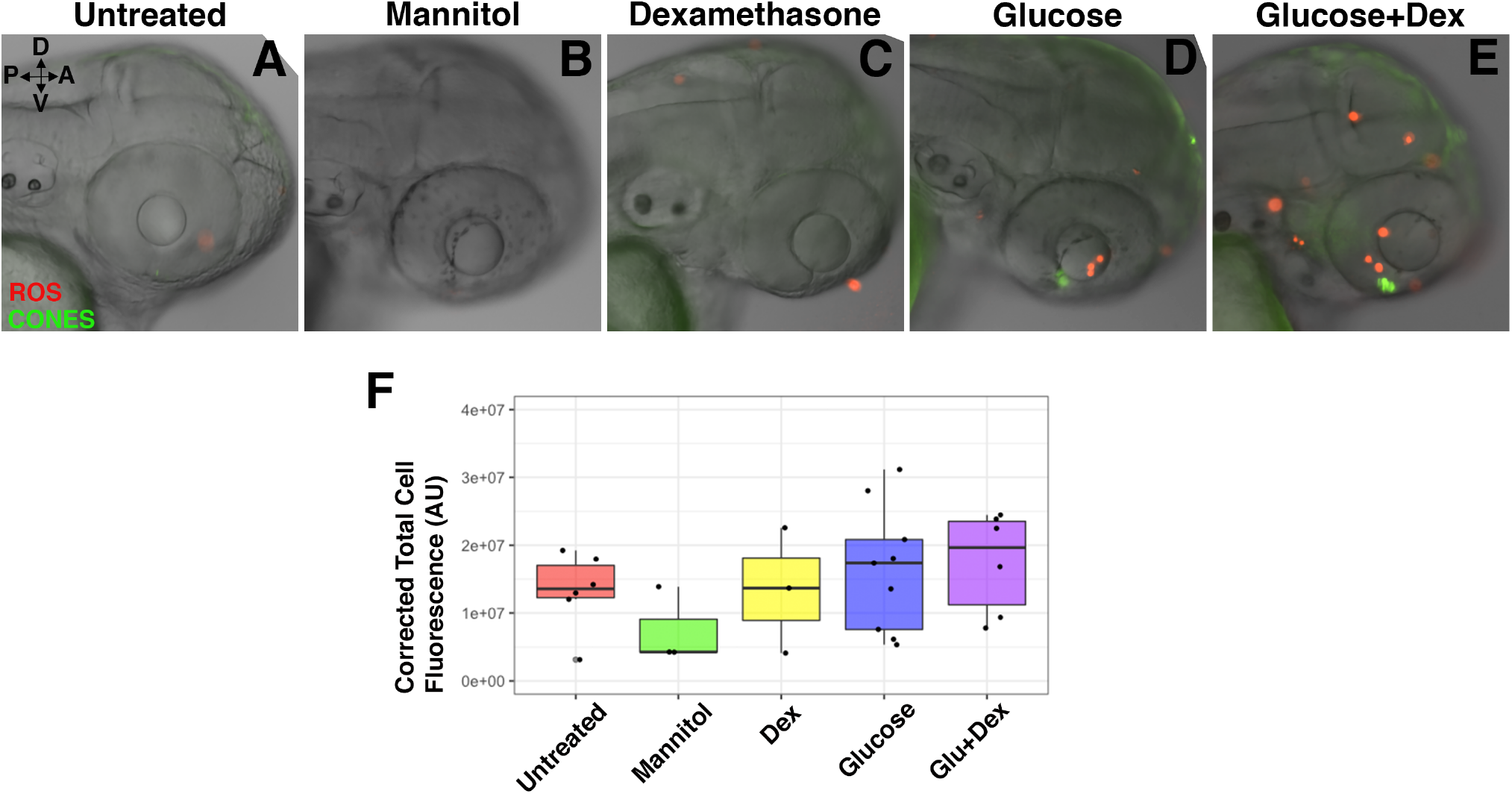
Increased superoxide production in hyperglycemic larvae at 48hpf. Treated larvae were submerged in a superoxide probe and imaged with by fluorescence microscopy. An increase in superoxide production was observed throughout hyperglycemic larval heads (F), particularly within the eye (D, E), compared to controls (A, B, C). D, Dorsal; V, Ventral; A, Anterior; P; Posterior.

To get a better idea of the metabolic factors specifically impacted by embryonic hyperglycemia, we examined whether expression of enzymes involved in glucose metabolism was altered using a glucose probe array at 5 dpf. Interestingly, we found that expression of key glucose metabolism enzymes connected to ROS production, such as succinate dehydrogenase, glucose-6-phosphatase, and nitric oxide synthase, was upregulated in the heads of glucose and glucose+dex treated larvae compared to untreated controls (Fig. S6K). The dysregulation of these enzymes can have detrimental effects on cellular metabolism, particularly in the context of hyperglycemia.

### Embryonic hyperglycemia perturbs the timing of retinal cell differentiation

We next sought to understand whether hyperglycemia was affecting the cell cycle and differentiation of retinal cells during development. Previous work has suggested embryonic hyperglycemia suppresses the cell cycle via altered expression of Cyclin D1 and p21, reducing cell proliferation and differentiation [38]. As mentioned above, ROS production has also been shown to regulate cell cycle exit in the developing retina [35]. To investigate the timing of retinal progenitor differentiation in our two models of developmental hyperglycemia, we tracked the fate of cells which were proliferating from 48-50 hpf. During this window of retinal development, photoreceptor progenitors are beginning to exit the cell cycle and differentiate. Ganglion and amacrine cells have mostly differentiated before 48 hpf, leaving primarily early photoreceptors and later neurons of the inner nuclear layer to differentiate in the targeted time window [39]. We exposed developing control and hyperglycemic embryos to EdU from 48-50 hpf and collected larvae at 5 dpf to visualize the fate of the EdU positive cells based on their location in the mature retina. We observed that the majority of EdU positive cells were located in the inner nuclear layer across all treatments and genotypes, which aligns with known developmental timing (Fig. 7F and J). For untreated, mannitol treated, and wildtype larvae, there was also a large proportion of EdU positive cells among the photoreceptor nuclei of the outer nuclear layer, indicating that, as expected, many of the proliferating RPCs at 48-50 hpf ultimately differentiated into photoreceptors. Very few EdU+ cells were in the ganglion cell layer at 5 dpf (Fig. 7A-B, F), consistent with the majority of retinal ganglion cells having already exited the cell cycle at the time of EdU exposure. In contrast, in hyperglycemic retinas, the second highest concentration of EdU positive cells was observed in the ganglion cell layer (Fig. 7 C-D, F), which suggests that unlike the controls, retinal ganglion cells had not yet exited the cell cycle at the time of EdU administration. Moreover, there were significantly fewer EdU positive cells in the outer nuclear layer (ONL) of hyperglycemic retinas at 5 dpf compared to controls, and the intensity of the EdU signal was also lower in cells of the ONL, suggesting that these cells were derived from retinal progenitors that had gone through more cell divisions than their counterparts in control retinas. Intriguingly, the total number of EdU+ cells was significantly increased in *pdx1* mutants compared to their wildtype siblings, but this was not observed in glucose or glucose+dex treated larvae (Fig. 7E vs. 7I). This is the first phenotype to be strikingly different between the genetic and nutritional model, suggesting that there could be a differential role of insulin in retinal progenitor cell (RPC) proliferation and differentiation. Taken together, these data indicate that embryonic hyperglycemia causes a delay in cell cycle exit and differentiation of RPCs. Since we did not observe a difference in ganglion, amacrine, and bipolar cell number between control and hyperglycemic retinas at 5 dpf, these cell types must “catch up” to normoglycemic levels of differentiation by 5 dpf. However, hyperglycemia appears to have prolonged impact on the number of retinal photoreceptors and horizontal cells.

**Fig. 7:**
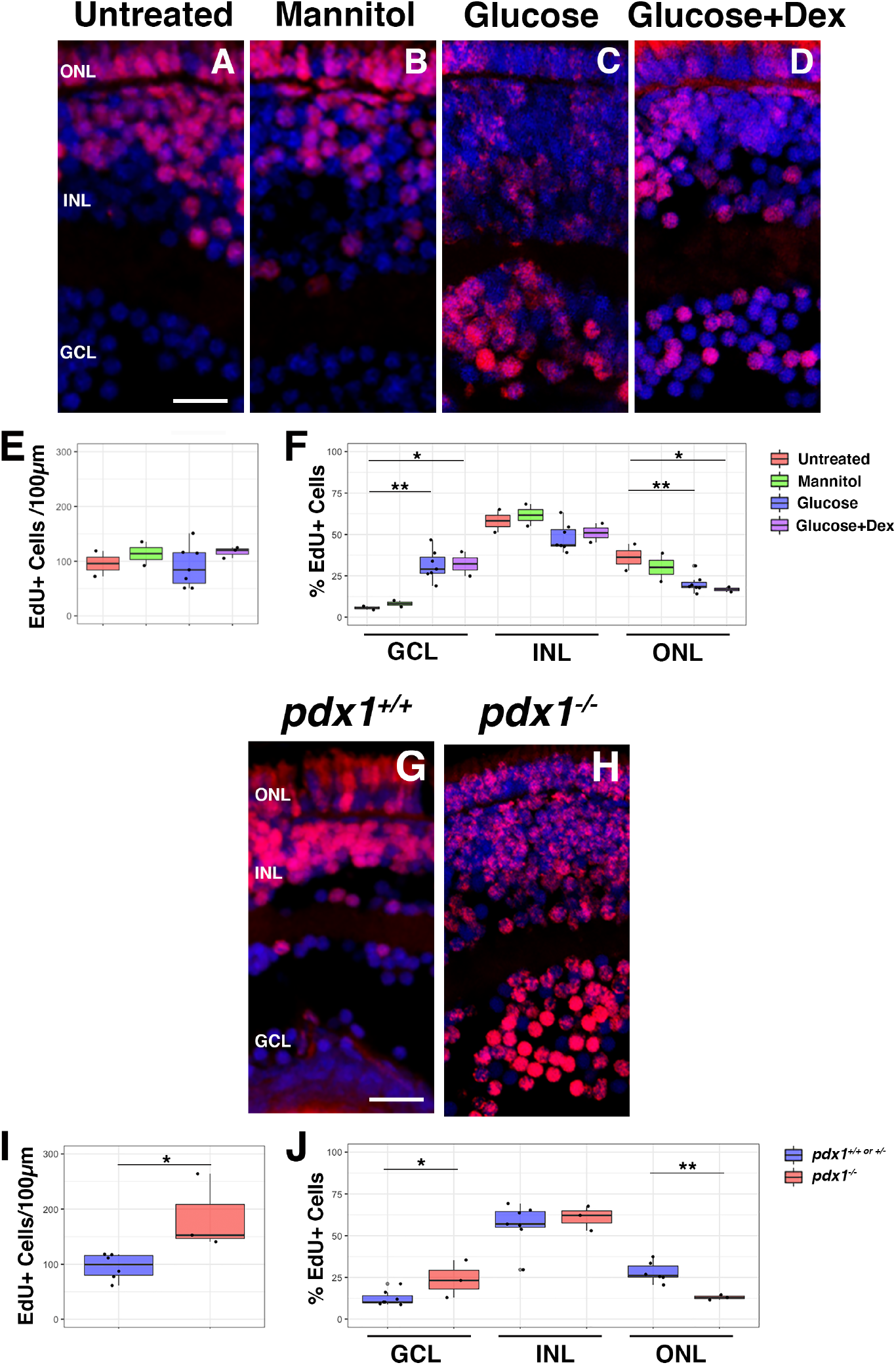
Hyperglycemic larvae exhibit increased retinal progenitor cell proliferation and delayed differentiation. Hyperglycemic embryos were exposed to EdU from 48-50 hpf then raised to 5 dpf and the cryosectioned. Glucose and Glucose+Dex treated larvae exhibited a significant increase in the percent of EdU+ cells in the GCL and significant decrease in the percent of EdU+ cells in the ONL (C, D, F) compared to untreated and mannitol treated larvae (A, B). The total number of EdU+ cells per 100µm did not significantly differ across treatments (E). Pdx1 mutant larvae displayed a significant increase in the number of EdU+ cells per 100 µm compared to wildtype (I). The distribution of EdU+ cells also showed a significant increase in the GCL and decrease in the ONL in the mutants (H) compared to wildtype (G, J). Scale bars: 10µm. * indicates p<0.05; ** indicates p<0.01

To determine whether reducing ROS could rescue photoreceptor number in hyperglycemic retinas, we treated the nutritional model and *pdx1* embryos with superoxide dismutase, but had difficulty establishing a treatment concentration which did not induce global side effects (data not shown). Therefore, we treated hyperglycemic embryos with the antioxidant methylene blue [40]. Methylene blue co-treatment produced an 18% increase in cone photoreceptors in glucose treated as well as glucose+dex treated larvae compared to no co-treatment (Fig. 8E). The increase in photoreceptors due to methylene blue supports our hypothesis that ROS production contributes to the decrease in photoreceptors in hyperglycemic retinas.

**Fig. 8:**
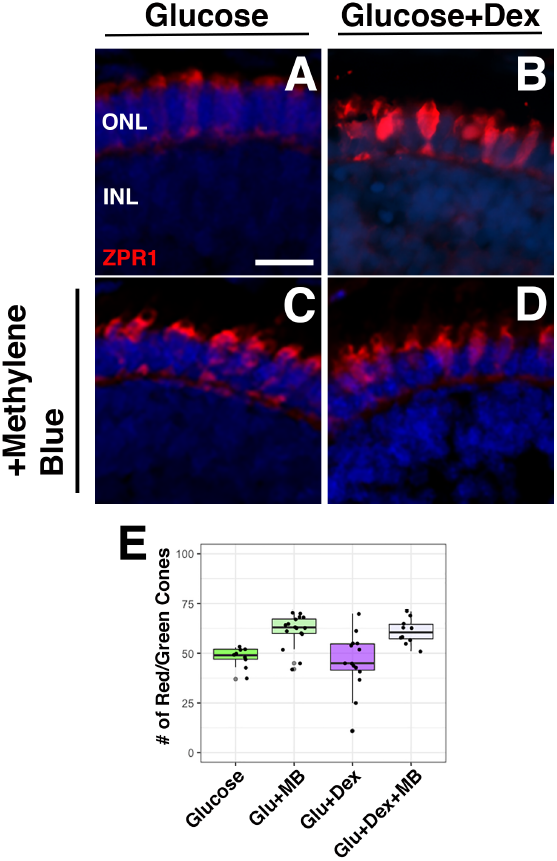
Co-incubation of methylene blue with nutritional model shows increased photoreceptors in hyperglycemic larvae. Nutritional model embryos were co-treated with methylene blue. Glucose and Glucose+Dex treated embryos showed an increase in red/green cone photoreceptors (C, D) following treatment with methylene blue at 5dpf (E) compared to no methylene blue treatment. Scale bar: 10µm. * indicates p<0.05

### Larvae that experienced embryonic hyperglycemia show a persistent decrease in cone photoreceptors after return to normoglycemia

The EdU pulse-chase experiment revealed a delay in retinal cell type differentiation which resulted in a decrease of photoreceptors at 5 dpf. To evaluate whether or not larvae could recover to produce photoreceptors at equivalent numbers to controls, we placed hyperglycemic larvae back into normal fish water at 5 dpf and collected them two days later at 7 dpf. We examined retinal sections by immunohistochemistry and found that cone photoreceptors were still significantly reduced in glucose and glucose+dex treated larvae compared to controls (Fig. S7I) while the number of rod photoreceptors across treatments was not significantly different (Fig. S7J). The persistent reduction in cones is indicative of potential long-term consequences of hyperglycemia on retinal cell maintenance and color vision. This is particularly important considering the susceptibility of cone photoreceptors to damage in adult models of hyperglycemia.

## Discussion

Diabetic retinopathy is a prevalent complication of diabetes and a principal cause for acquired blindness in adults [41]. A growing body of research has demonstrated that the degeneration of photoreceptor cells actually precedes the retinal vasculature defects in diabetic retinopathy [42]. This underscores that the photoreceptors are a critical focal point for initiating retinal pathology as a result of chronic hyperglycemia. As the population of people with diabetes rapidly grows, research efforts must also be focused on studying the consequences of diabetes for pregnant mothers and the retinal development of their offspring.

While there is abundant research on heart development in hyperglycemic embryos, little work has focused on the retina specifically. In humans, one study showed that offspring of diabetic pregnancies had significantly thinner inner and outer macula as well as lower macular volume [43]. Considering the macula contains the highest density of cone photoreceptors in the human eye, this study suggests a potential deleterious effect of diabetic pregnancy on retinal cone photoreceptor development; however, there is a need to visualize the effects on photoreceptor number and morphology in more detail. Our study can fill this gap by combining whole mount imaging and retinal sections to visualize, quantify, and assess the morphology of retinal cells that have developed under hyperglycemic conditions. This is the first study to characterize the development of photoreceptors specifically across two models of embryonic hyperglycemia. One of the most striking findings was the decrease in photoreceptor number in hyperglycemic larvae, with the remaining photoreceptors displaying stunted outer segments – these findings were observed in both nutritionally and genetically induced hyperglycemic models. Without full outer segments, we hypothesized the hyperglycemic larvae could have visual defects, which was supported by their poor optokinetic response.

Our results demonstrate how embryonic hyperglycemia affects the retina short term, but we were also interested in long term effects. Looking at retinal sections of treated embryos from the nutritional model at 7dpf, we found that cone photoreceptors continued to be reduced in number even after the larvae were returned to normoglycemic conditions, whereas rod photoreceptor number seemed to catch up to those of control larvae. This indicates an issue in cone photoreceptor programming, metabolism, and/or maintenance which may result in longer-term visual defects as well as potential susceptibility to degeneration. Considering cone photoreceptor degeneration is an early consequence of persistent hyperglycemia in adult zebrafish, it will be important to study how cone photoreceptors in zebrafish that experienced embryonic hyperglycemia respond to a “second hit” of hyperglycemia as adults. Further, understanding how embryonic hyperglycemia affects the retina both short and long term is imperative to identifying and timing therapeutics during development to prevent lasting vision problems.

In addition to photoreceptor phenotypes, a recent study utilizing zebrafish to study retinal development showed that exposure to high levels of glucose causes a decrease in retinal ganglion cells and Müller glia as well as an increase in infiltrating macrophages [16]. That study utilized a different timecourse of submersion and a much higher glucose concentration than our study. These differences in methodology may explain our discrepant findings in Müller glia cell number, with our model showing not only a slightly higher number of Müller glia in hyperglycemic zebrafish, but also an abnormal morphology suggestive of reactive gliosis. We also did not observe a reduction in retinal ganglion cell number in either model, which could be due to the lower glucose concentration used, or to different methods to visualize the ganglion cells.

Given the abnormal Müller glia morphology and the elevated cell death in the inner nuclear layer of hyperglycemic retinas, it is possible that reactive gliosis is a consequence of the decrease in photoreceptors or the altered metabolic environment, or a combination of both considering the role and needs of Müller glia in the retina. Interestingly, the only other class of retinal neuron that showed differing numbers or morphology in our study were the horizontal cells, which were reduced in *pdx1* mutants, glucose treated, and glucose+dex treated larvae. The reduction in horizontal cells may be directly related to the reduction in photoreceptors, as previous studies have shown that horizontal cells play a role in cone/rod distribution and synaptogenesis [44].

Considering the link between hyperglycemia, ROS production, and cell death, we hypothesized that the decrease in photoreceptors could be due to apoptosis. While we did observe an increase in apoptotic cells in hyperglycemic larvae, the TUNEL+ cells were not localized to the outer nuclear layer where photoreceptors reside. Rather, most of the TUNEL+ cells were found in the inner nuclear layer, with many co-localizing with Müller glia nuclei. This could indicate that Müller glia phagocytose dying photoreceptors in hyperglycemic retinas, as has been observed in other models of photoreceptor degeneration [45]–[47] and may also account for the apparent gliotic response. However, because the overall number of TUNEL+ cells was relatively low at all time points, it is likely that additional factors contribute to the decrease in photoreceptor number in response to hyperglycemia, such as altered photoreceptor cell differentiation.

A recent study by Albadri et. al., showed retinal cell developmental delay can be induced by lipid peroxidation products such as 4-HNE, which downregulate HDAC1 to keep cells in a proliferative state, preventing differentiation [48]. Production of ROS such as H_2_O_2_ is an upstream event leading to lipid peroxidation, and in human endothelial cell lines modeling hyperglycemia [49]. The link between photoreceptors and ROS production has yet to be elucidated in terms of mechanism. Given our observation of increased ROS in hyperglycemic retinas at 48 hpf and 5 dpf, we hypothesize that genetic or nutritionally induced embryonic hyperglycemia results in increased ROS production which may cause retinal cell differentiation delay, and abnormal photoreceptor morphology.

Indeed, utilizing EdU birth-dating, we found that hyperglycemic larvae exhibited a striking deviation from the expected timing of RPC cell cycle exit during what should have been the window of photoreceptor differentiation. The significant increase in EdU+ cells in the retinal ganglion cell layer indicates a lag in retinal cell type differentiation in hyperglycemic larvae that is likely a major contributor to the decreased number of photoreceptors. In *pdx1* mutants, we also found a significant increase in the overall number of EdU+ cells that was not observed in our nutritional model, which indicates loss of *pdx1* specifically results in an increase in RPC proliferation. Whether this is related to the lack of insulin producing cells in *pdx1* mutants warrants further investigation. Importantly, while RGCs and other retinal neurons eventually overcome this delay and differentiate in comparable numbers to control retinas at 5 dpf, cone photoreceptors remain reduced relative to controls even after an additional period of normoglycemic conditions. This result suggests that while most retinal neurons exhibit adaptive plasticity in their developmental timing, cones may be particularly sensitive to early metabolic derangement. Taken together, our data now connect embryonic hyperglycemia with increased ROS production, increased RPC proliferation, and extended perturbations to photoreceptor differentiation.

Given our ROS data in the nutritional model, we tested whether an antioxidant, methylene blue [40], could rescue the decrease in photoreceptors in hyperglycemic larvae. We found that addition of methylene blue in our nutritional model increased the number of cone photoreceptors, with morphology which better resembled controls, particularly the outer segments (Fig. 8). It is unclear though whether methylene blue improved the overall health of the larvae or specifically rescued retinal phenotypes. We hypothesize that a combination of antioxidants and ROS inhibitors with a more targeted delivery method could be necessary to rescue the decrease in photoreceptors specifically. Further, analysis of the particular forms of ROS as well as other reactive compounds that are produced in response to hyperglycemia is needed to identify better targets for pharmacologic intervention.

To better understand metabolic factors that are dysregulated and may link hyperglycemia with ROS production, we used a glucose probe array to quantify expression of enzymes related to glucose metabolism. Under normoglycemic conditions, photoreceptors take up glucose which is broken down and converted to lactate via aerobic glycolysis [50]. In a hyperglycemic state, there is an influx of glucose undergoing glycolysis which can alter the expression of various enzymes in the glycolytic pathway, that can have further downstream effects [51]. Additionally, lactate functions as a nutrient source for Müller glia and feeds the RPE to suppress glycolysis, in turn increasing glucose transport to photoreceptors [52]. Downstream metabolic processes such as the Kreb’s Cycle and Electron Transport Chain (ETC) are affected by this as well due to their utilization of metabolic intermediates [53]. We found a significant increase in enzymes that play important roles in NADH production that feeds into the ETC where it is oxidized at complex I and reacts with reduced flavin. Under normal conditions, an electron can be passed from flavin to O_2_ which forms O_2_^-^ (superoxide) during NADH oxidation/flavin reduction [54]. When excess NADH is fed into the ETC, superoxide production is increased, which we noted in our hyperglycemic larvae. Superoxide can induce DNA damage [55] and delays in cell differentiation. ROS production can be further perpetuated via methylglyoxal [56] which is upregulated in hyperglycemic states [57]. Studies have shown increased methylglyoxal induces vasculature damage, activation of glial cells, and disrupts retinal function in the presence and absence of hyperglycemia [58]. Given our results, we created a model that connects the metabolic effects of hyperglycemia to the photoreceptor phenotypes we observed (Fig. 9). We propose that hyperglycemia induces delayed photoreceptor differentiation early in retinal development while a gliotic response is induced later via ROS production and cell death. It will be important for future studies to look closely at the highlighted enzymes of our model in terms of pathways which they affect and in coordination with one another to better understand key components in the glycolytic pathways that would serve as promising therapeutic targets.

**Fig. 9:**
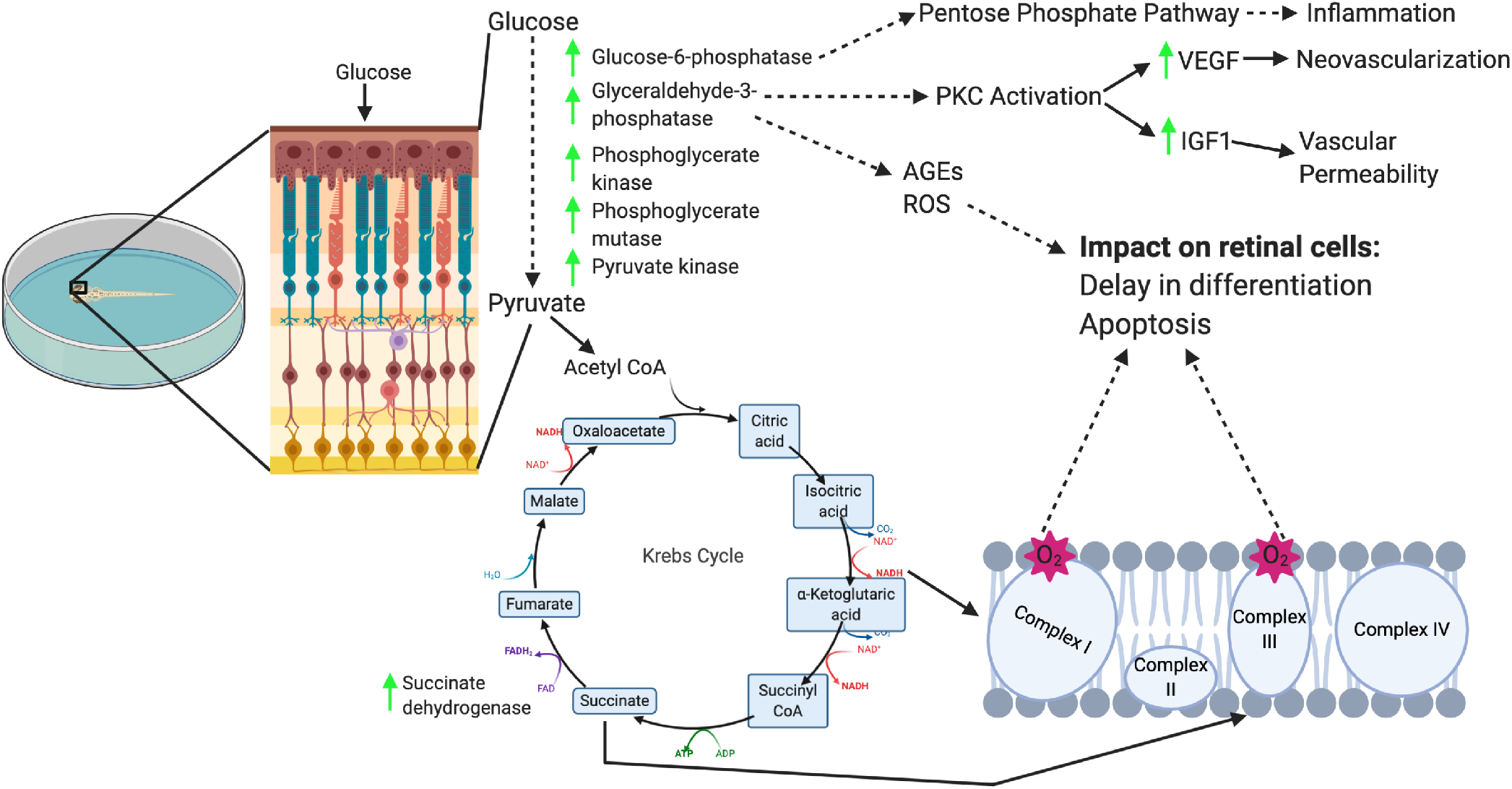
Proposed model for how hyperglycemia alters retinal development. When hyperglycemia is induced, there is an increase of glucose flux to photoreceptors. The increase in glucose consumption by photoreceptors increases the rate of glycolysis, thereby increasing expression of enzymes related to this pathway including glucose-6-phosphatase, glyceraldehyde-3-phosphatase, phosphoglycerate kinase, phosphoglycerate mutase, and pyruvate kinase, as supported by the glucose probe array data. Upregulation of glucose-6-phosphatase in particular is indicative of potential pentose phosphate pathway activation which can induce inflammation and mediate ROS production. Glyceraldehyde-3-phosphatase upregulation may lead to PKC activation which increases VEGF and IGF-1 expression to induce neovascularization and vascular permeability, respectively. AGEs and ROS production has also been linked to glyceraldehyde-3-phosphatase upregulation which can cause DNA damage, PARP activation, and apoptosis in retinal cells. At the tail end of glycolysis, pyruvate feeds into the Tricarboxylic Acid (TCA) Cycle, producing NADH via reduction from NAD to convert isocitrate to alpha-ketoglutarate as well as succinate dehydrogenase. NADH feeds into the electron transport chain (ETC) at complex I and succinate dehydrogenase feeds into complex III reacts with reduced flavin. When excess NADH is fed into the ETC, superoxide production is increased, which can induce DNA strand breaks, cell death, and potential disruption in retinal cell differentiation.

Embryonic development requires a complex coordination of events at the cellular level which can be altered through metabolic stressors. As hyperglycemia becomes increasingly common, it is necessary to understand how it affects development of all tissues. With the retina being a major consumer of glucose and given the known deleterious effects of chronic hyperglycemia, examining retinal development in a hyperglycemic state is critical. Our findings clearly show that hyperglycemia negatively impacts retinal development, and suggest a potential mechanism of action, which may inform the search for therapeutic targets. Our future studies will include elaborating on our embryonic hyperglycemia models through metabolic assays and alternative “rescue” experiments. We also look forward to studying how the hyperglycemic larval retina continues to develop, maintain itself, and function long term.

## Materials and Methods

### Zebrafish Lines and Maintenance

Zebrafish were bred, raised, and maintained in accordance with established protocols for zebrafish husbandry. All zebrafish lines were bred and raised at 28.5°C on a 14-hour light:10-hour dark cycle as previously described [59]. The Tg(3.2TαC:EGFP) transgenic line (TαC:EGFP), previously described [26], was generously provided by Susan Brockerhoff (University of Washington, Seattle, WA). The Tg(XlRho:EGFP) transgenic line (XOPS:GFP), has been previously described, and was obtained from James Fadool (Florida State University, Tallahassee, FL, USA; Fadool, 2003). The Tg(gfap: GFP)mi2001 or GFAP:GFP transgenic line [60], was obtained from the Zebrafish International Resource Center (ZIRC, Eugene, OR). Heterozygous *pdx1*^*sa280*^ adults were generously provided by Tim Mulligan (Johns Hopkins University, Baltimore, MD). *Pdx1* genotype was determined by PCR using the primers 5’-TGGCTCATGTGCTCGTGTA-3’ and 5’-GTGCGTGTGAGATTTGGTTG-3’ followed by RFLP analysis with DraI. Embryos were anesthetized with Ethyl 3-aminobenzoate methanesulfonate salt (MS-222, Tricaine; Sigma-Aldrich Corp., St. Louis, MO). All animal procedures were carried out in accordance with guidelines established by the University of Kentucky Institutional Animal Care and Use Committee and the ARVO statement on the use of animals in research.

### Generating hyperglycemic zebrafish embryos

GFAP:GFP, XOPS:GFP and TαC:eGFP adult fish were in-crossed to generate embryos that were sacrificed at the following time points: 48 hpf, 72 hpf, 96 hpf, and 5 dpf. To generate hyperglycemic embryos/larvae, 10 hpf embryos were dechorionated with pronase (Sigma), and randomly sorted into groups of 25 and placed in the following treatments: untreated fish water, 50 mM glucose, 50 mM mannitol, 10μM dexamethasone (Sigma), or 50 mM glucose + 10 μM dexamethasone (glucose+dex) in 1x Phenylthiourea (Sigma) dissolved in fish water. At each time point of interest, heads were removed for cryosectioning, mRNA, or protein extraction, while the body was used to quantify whole body glucose concentration. For the *pdx1* line, larvae were trisected: heads were used for retinal sectioning, mid-body area was used to quantify glucose using the glucose assay, and genomic DNA was extracted from the tail for genotyping.

### Glucose concentration quantification

A glucose colorimetric assay kit from Biovision (Biovision, Milpita, CA) was used to quantify whole body glucose concentration, following the manufacturer’s instructions. In short, following sacrifice, animals were homogenized in glucose assay buffer, mixed with a glucose oxidation enzyme and a colorimetric probe, and incubated for 20 minutes at 37°C. A spectrophotometer was used to quantify glucose concentration. Reads were translated into pmol/larvae through generation of a standard curve from a set of samples of defined glucose concentrations.

### Cryosections and cell counts

Embryos were fixed overnight in 4% paraformaldehyde, then incubated overnight in 10% followed by 30% sucrose at 4°C. Transverse 10µm sections were taken beginning in the anterior tip of the head, moving posteriorly through the eye. For imaging and cell quantification, only sections containing an optic nerve were used for consistency. Photoreceptors in the dorsal, central, and ventral portions of the retina were quantified and normalized to the curvilinear length of the outer nuclear layer. For the HuC/D, PKCα, and Prox1 quantification, counts were conducted on 50µm wide regions of interest, 50µm dorsal to the optic nerve for consistency. Images were taken using a 20x objective on a Leica SP8 Confocal microscope or a Nikon Eclipse inverted fluorescent microscope (Eclipse Ti-U, Nikon Instruments). At least 10 embryos were analyzed per treatment/genotype, and 3 separate biological replicates were performed for each experiment. Statistics were conducted using two-factor, unpaired t-test using GraphPad software. P-values less than 0.05 were considered significant and are indicated by *, p< 0.01 is indicated by **, and p<0.001 by ***. Boxplots were generated using R (version 3.6.2)/R studio (version 1.2.5033) ggplot2 package [61]. Figures 2 and 9 were created using Biorender (biorender.com). All figures were constructed using Photoshop (Adobe version 21.0.2).

### Immunohistochemistry, TUNEL and EdU Assay

Sectioning and immunohistochemistry were conducted as previously described [62] and immunolabeled sections were imaged on either a Nikon inverted (Nikon Ti-U) or confocal microscope (Leica SP8, Leica). The following antibodies were used: anti-Zpr3 (photoreceptor outer segments, mouse, 1:100, ZIRC), anti-Huc/D (ganglion and amacrine cells, mouse, 1:40, ZIRC), anti-PKCα (bipolar cells, mouse, 1:100, Santa Cruz Biotechnology, Dallas, TX), and anti-Prox1 (horizontal cells, rabbit, 1:1000, Acris, San Diego, CA). Alexa fluor conjugated secondary antibodies (Invitrogen, Grand Island, NY) and Cy-conjugated secondary antibodies (Jackson ImmunoResearch, West Grove, PA) were used at 1:200 dilution. Slides were incubated in 4’,6-diamidino-2-phenylindole (DAPI) to label nuclei (1:10,000 dilution, Sigma). TUNEL assay was conducted with ApopTag Fluorescin Direct In Situ Apoptosis Detection Kit (Millipore, Billerica, MA) on retinal cryosections according to manufacturer’s instructions. EdU assay was conducted with the EdU Cell Proliferation Assay Kit (EdU-555, Millipore, Billerica, MA). Embryos were soaked in 0.75mM EdU for 2 hours at room temperature from 48-50 hpf, then washed in fresh fish water, and raised to 5 dpf. Heads were cryosectioned and the Clik-It assay was conducted on retinal cryosections according to the manufacturer’s instructions.

### Metabolic pathway analysis

A Zebrafish Glucose Metabolism RT2 Profiler PCR Array (Qiagen, location) was used to quantify 86 different enzymes involved in pathways related to glucose metabolism (glycolysis, TCA cycle, electron transport chain, etc.) according to the manufacturer’s instructions. Briefly, mRNA was extracted, and cDNA generated as previously described [59], from larval heads at 5 dpf for all treatment groups and *pdx1* genotypes. cDNA, SYBR Green, and ultra-purified water were mixed and pipetted into the plate and a Roche Light Cycler (Roche, city) was used to run a program consisting of 45 cycles, 95°C for 15 seconds, 60°C anneal for 60 seconds. Gene expression was quantified with Roche Light Cycler Analysis program and normalized to the housekeeping gene, beta-actin.

### ROS production visualization

Treated and control 5 dpf larvae were submerged in 5mM MitoSOX ROS probe (Invitrogen, Carlsbad, CA) for 20 minutes at 28°C in the dark. Larvae were washed 3 times in fish water, mounted in 4% low melting agarose/1% tricaine, and imaged with a Leica SP8 confocal microscope. Following imaging, eyes were microdissected for retinal sections and imaged on the Leica SP8 Confocal microscope. ROS production was quantified by fluorescent pixel density using ImageJ.

### Optokinetic Response Test

Optokinetic Response Test was conducted as described by [63]. Larvae were assessed at 5 dpf, 1 at a time for 1 minute at 6 rotations per minute. Each larva was tested 4 times, twice in each direction of the rotating drum, and averaged. Saccadic eye movements were quantified and compared across treatments and genotypes with an ANOVA/unpaired t-test.

## End Matter

### Author Contributions and Notes

K.F.T-T. and A.C.M. conceived and designed the experiments and wrote the manuscript. K.F.T-T. performed the experiments. K.F.T-T. and A.C.M. prepared the figures. All authors reviewed the manuscript.

## Acknowledgments

The authors would like to thank Evelyn Turnbaugh and Lucas Vieira Francisco for exceptional zebrafish care. We also thank Dr. Hannah Henson for conceptual input and Dr. Cagney Coomer for editorial input. This work was supported by grants from the NIH National Eye Institute (R01EY021769, to A.C.M.), the WUSTL Diabetes Research Center, an NIH-funded program (P30DK020579) supported by the NIDDK (to A.C.M.), the National Science Foundation Bridge to the Doctorate Fellowship (NSF HRD 2004710, to K.F.T-T.), and the University of Kentucky Lyman T. Johnson graduate fellowship (to K.F.T-T.).

