## Supplemental data for "Embryonic hyperglycemia perturbs the development of specific retinal cell types, including photoreceptors"

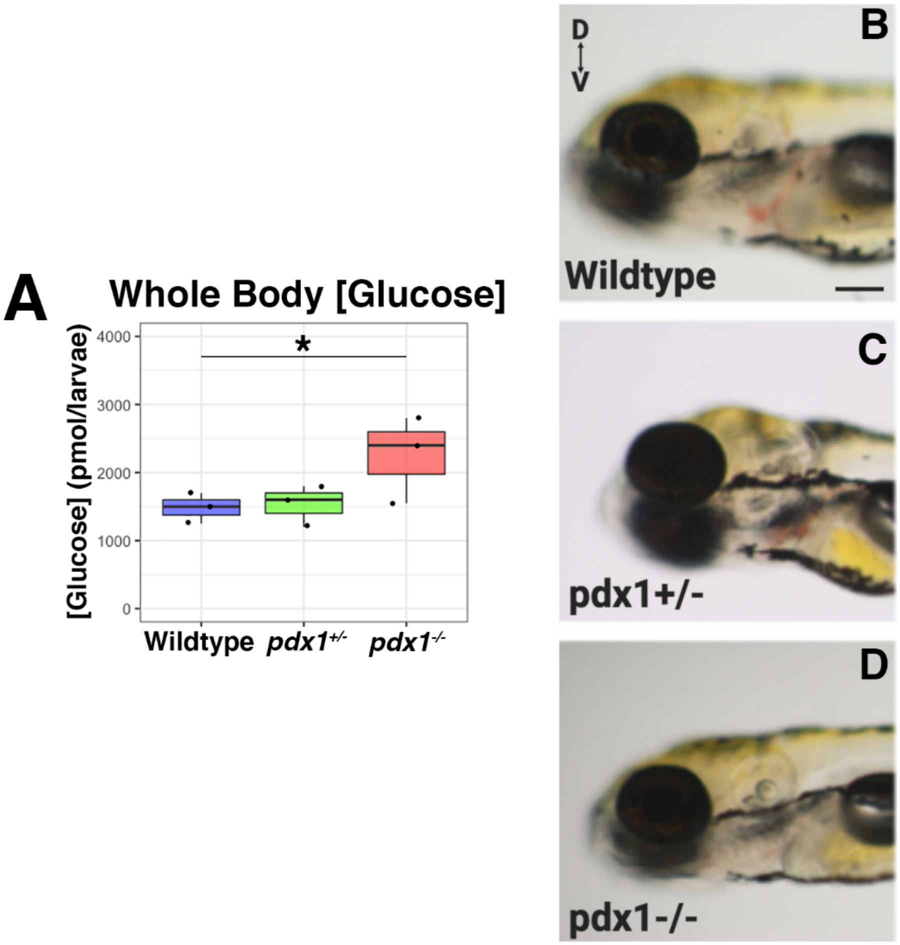


**Supplemental Figure 1: 5 dpf *pdx1* mutant larvae are hyperglycemic with normal overall morphology.** Progeny of *pdx1*$^{+/-}$ adults were sacrificed at 5 dpf. Genomic DNA was extracted from the remaining tail to genotype via PCR followed be RFLP. Whole body glucose was significantly elevated at 5 dpf (A). Images of whole larvae showed normal gross eye morphology and size across genotypes (B-D). Scale bar: 100μm. * indicates $p<0.05$

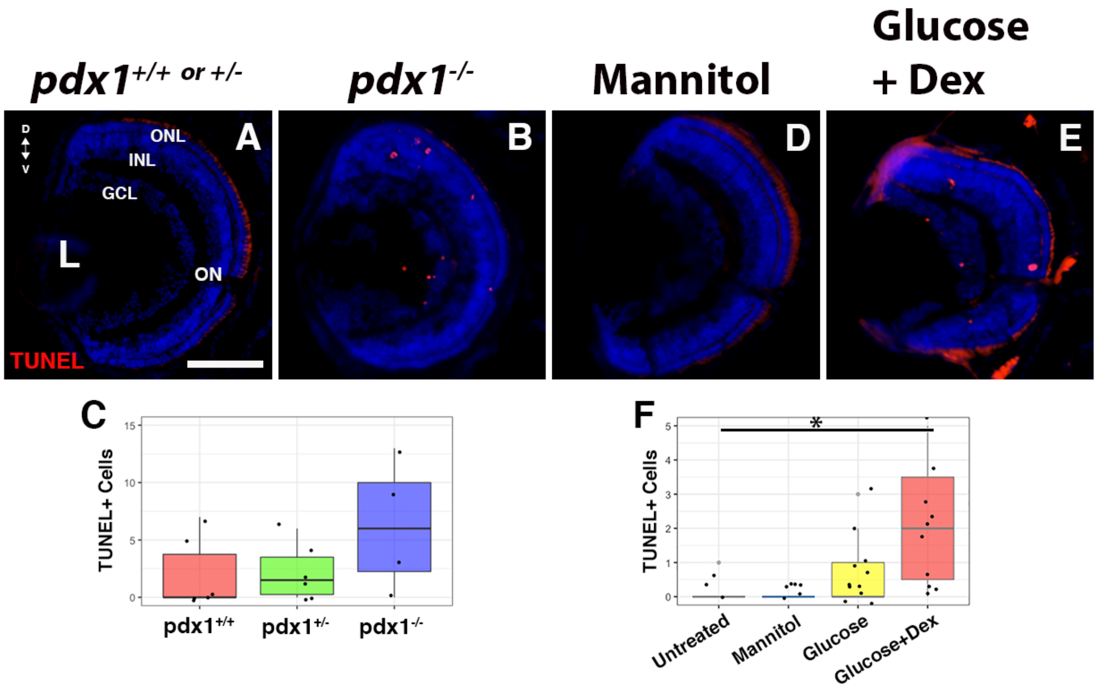


**Supplemental Figure 2: Apoptosis is increased in the retinas of hyperglycemic larvae.** Pdx1 larvae were genotyped, then both Pdx1 and treated larvae were sectioned at 5 dpf; a TUNEL assay was used to visualize apoptotic cells. The number of TUNEL+ cells was elevated in hyperglycemic larvae (B, E) compared to controls (A, D). Statistically significant elevation in apoptotic cells was observed in glucose+dex treated larvae (F). D, Dorsal; V, Ventral; ONL, Outer Nuclear Layer; INL, Inner Nuclear Layer; GCL, Ganglion Cell Layer; L, Lens. Scale bars: 50μm (A). * indicates $p<0.05$

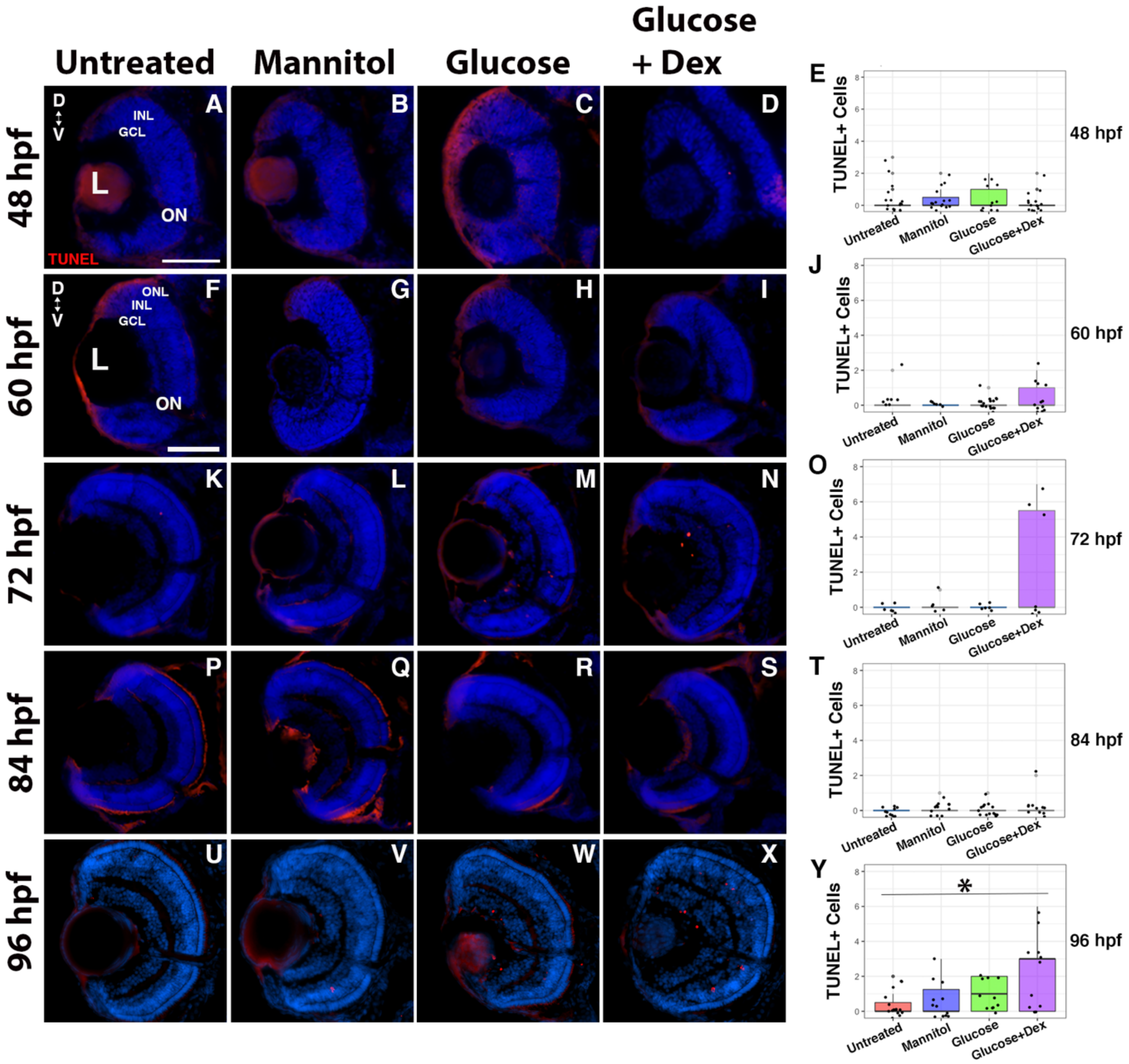


**Supplemental Figure 3: Cell death in the retina is elevated in hyperglycemic embryos throughout development**. Treated larvae were sectioned every 12 hours from 48 hpf to 5 dpf, and a TUNEL assay was conducted to visualize apoptotic cells. At 48, 60 and 72 hpf, apoptosis was observed in glucose and glucose+dex treated embryos only, primarily noted in the GCL and INL (F-J, L-O). Minimal apoptosis was noted at 84 hpf across all treatments (U), and a significant increase in apoptosis was observed at 96 hpf in glucose+dex treated embryos (Y). D, Dorsal; V, Ventral; ONL, Outer Nuclear Layer; INL, Inner Nuclear Layer; GCL, Ganglion Cell Layer. Scale bars: 50μm (A, F). * indicates $p<0.05$

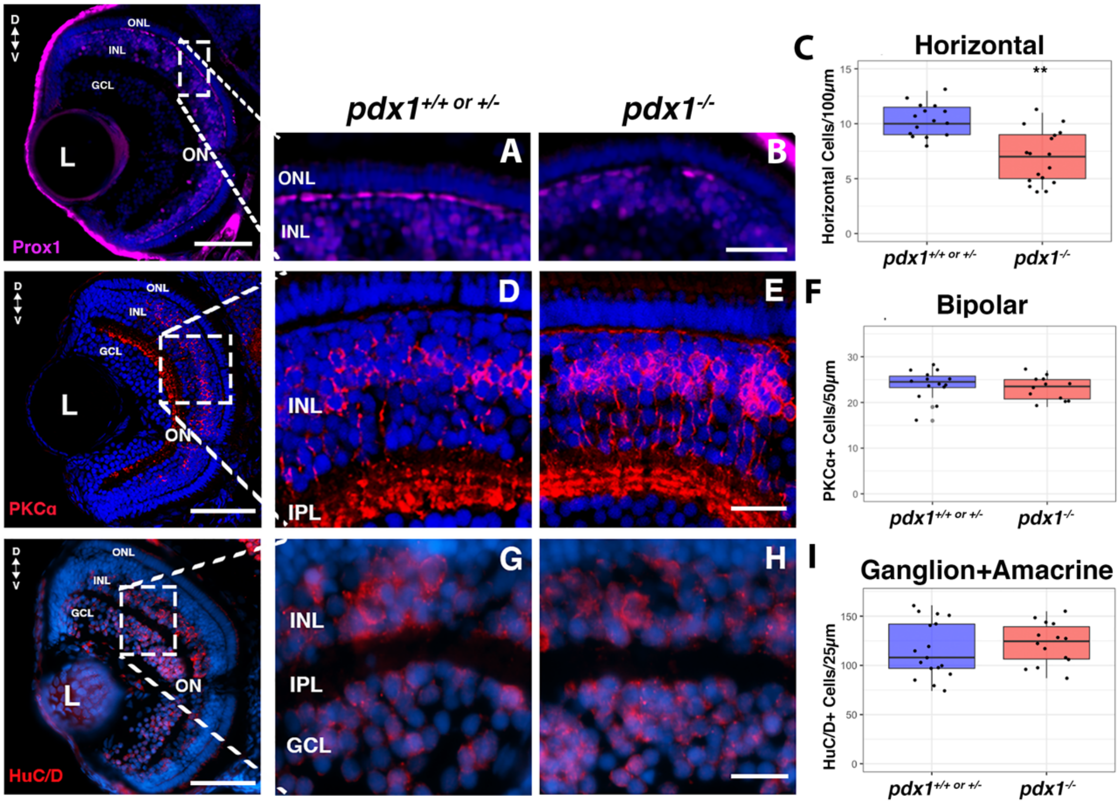


**Supplemental Figure 4: Horizontal cells are significantly reduced whereas ganglion, amacrine, and bipolar cells are unchanged in *pdx1*$^{-/-}$ larvae.** Pdx1 larvae were genotyped and sectioned at 5 dpf, then immunostained and imaged to visualize inner retinal neurons. Prox1 immunolabeling (A-B) revealed a significant reduction in the number of horizontal cells in *pdx1*$^{-/-}$ larvae (C). Bipolar (D-E), ganglion, and amacrine cell numbers (G-H) were not different across genotypes (F, I). White dashed boxes indicate region of retina where counts and analysis were conducted on each section. D, Dorsal; V, Ventral; ONL, Outer Nuclear Layer; INL, Inner Nuclear Layer; GCL, Ganglion Cell Layer; L, Lens. Scale bars: 100μM, 10μM (B), and 20μM. ** indicates $p<0.01$

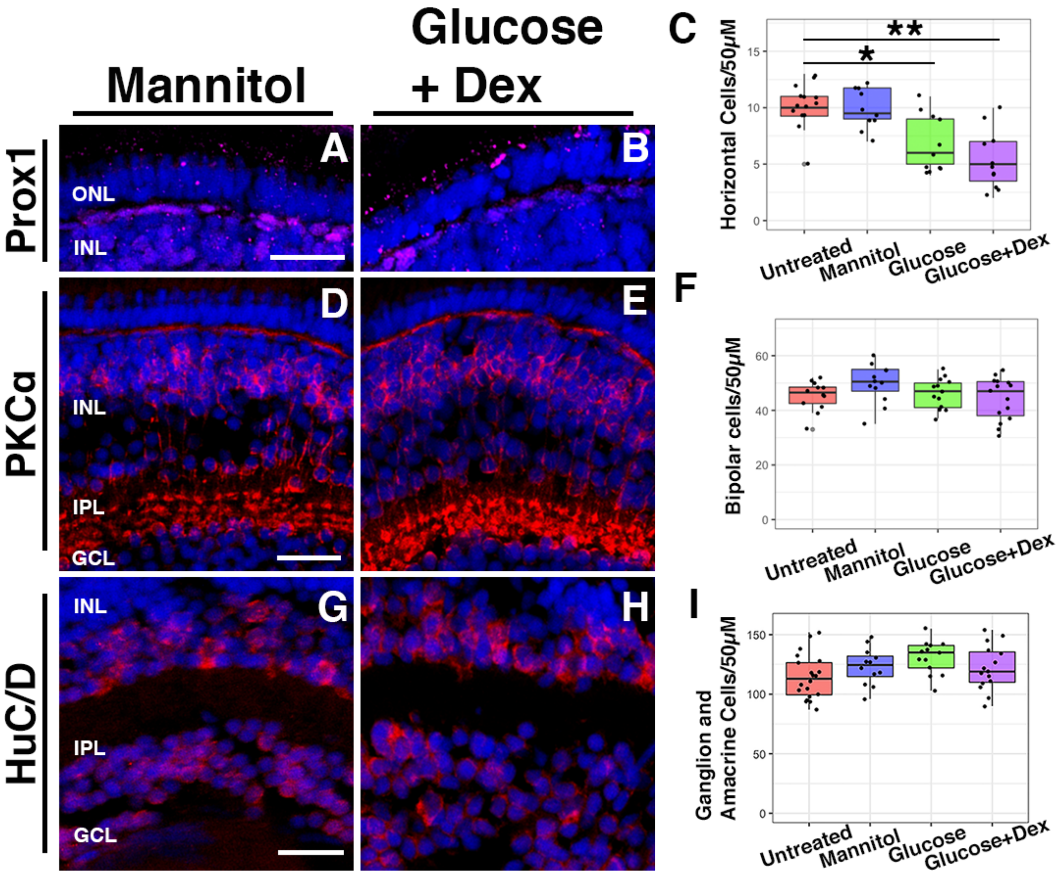


**Supplemental Figure 5: Inner retinal neurons are unchanged, while horizontal cells are reduced, in a nutritional model of hyperglycemia.** Treated larvae were sectioned at 5 dpf, then immunostained and imaged to visualize inner retinal neurons. Horizontal cells (A-B) were significantly reduced in glucose + dex larvae compared to controls (C). Bipolar cells showed no significant differences in number (F) or morphology (D-E). Ganglion and amacrine cell numbers (G-H) were also not different across treatments (I). White dashed boxes indicate region of retina where counts and analysis were conducted on each section. D, Dorsal; V, Ventral; ONL, Outer Nuclear Layer; INL, Inner Nuclear Layer; GCL, Ganglion Cell Layer; L, Lens. Scale bars: 10µm (A, G), and 20µm (D). * indicates $p<0.05$; ** indicates $p<0.01$

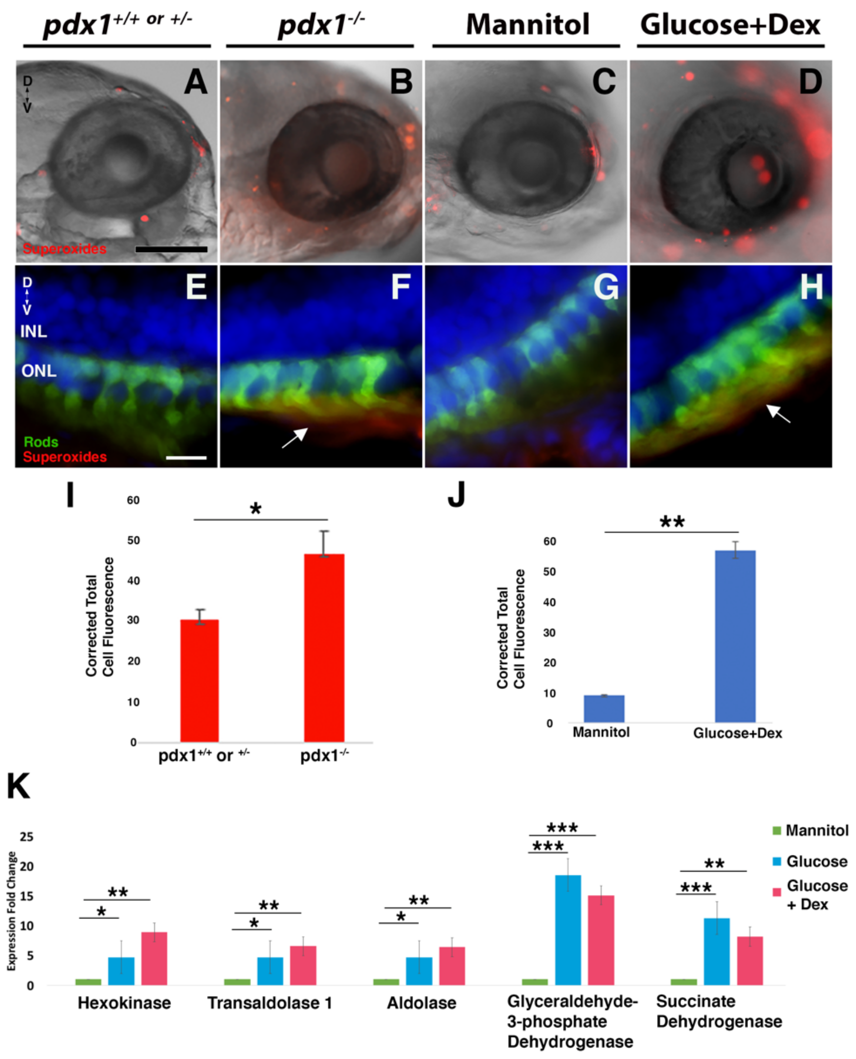


**Supplemental Figure 6: Increased superoxide production and abnormal glucose metabolism in 5dpf hyperglycemic larvae.** Pdx1 mutant and treated larvae were submerged in a superoxide probe and imaged with by fluorescence microscopy. A significant increase in superoxide production (I, J) was observed throughout hyperglycemic larval heads (B, D), particularly within the eye, compared to controls (A, C). Retinal sections of control and hyperglycemic XOPS:GFP larvae revealed colocalization of the superoxide probe among the outer segments of rod photoreceptors in hyperglycemic retinas (F, H; arrows). D, Dorsal; V, Ventral; ONL, Outer Nuclear Layer; INL, Inner Nuclear Layer. Scale bars: 100μm (A) and 10μm (E).  * indicates $p<0.05$; ** indicates $p<0.01$

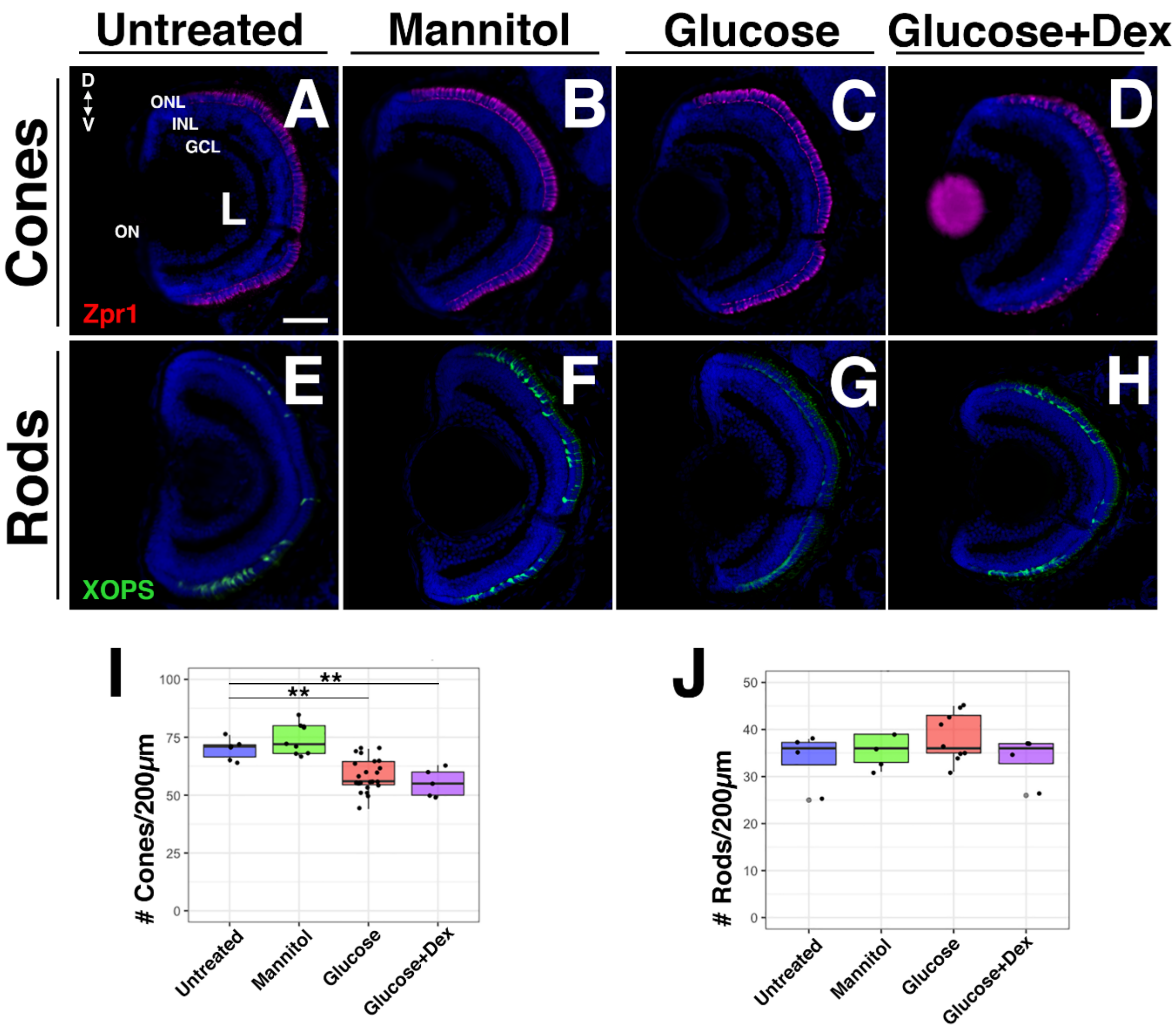


**Supplemental Figure 7: Cone photoreceptors are persistently reduced in juveniles that experienced embryonic hyperglycemia.** Treated larvae were placed in normal fish water at 5 dpf, then sectioned, immunostained, and imaged at 7 dpf to visualize photoreceptors. Zpr1 immunostaining revealed persistently reduced number of cone photoreceptors (I) in larvae that had previously experienced glucose + dex and glucose treatment (C, D) compared to untreated and mannitol treated (A, B). Rod photoreceptors showed no significant difference in number across treatments (J). D, Dorsal; V, Ventral; ONL, Outer Nuclear Layer; INL, Inner Nuclear Layer. Scale bars: 50µm (A, C, F). ** indicates $p<0.01$
